# Transformer-based framework uncovers state-dependent modular organization and conformational landscapes of the β-arrestin 1 C-terminal tail

**DOI:** 10.64898/2026.06.06.730629

**Authors:** Margaret Robinson, Van Ngo, Jonathan A. Javitch, Lei Shi

## Abstract

β-arrestins (βarr) regulate signaling and trafficking of G protein-coupled receptors (GPCRs) across diverse physiological and pathological processes. However, mechanistic understanding of how ligand-activated GPCRs engage and activate βarr remains limited, with the conformation of the entire βarr tail in the active state still unknown. Here, by comparatively analyzing temperature replica-exchange molecular dynamics simulation data of βarr1 in basal and active states, we investigated the conformational landscape of the βarr1 tail to elucidate its role in activation. To overcome limitation of conventional analyses in characterizing the vast conformational space sampled by the 62-residue tail, we developed a transformer-based autoencoder (TAE) framework that integrates attention-derived residue relationships and latent-space clustering to identify the tail’s modular organization and conformational substates, providing an interpretable description of how local residue interactions couple to large-scale conformational rearrangements. Using the conformationally constrained basal state as a control, we validated the framework by showing that it recovers interpretable conformational features. In the active state, the framework revealed a reorganized modular architecture, recovered key interactions identified through manual analysis, and uncovered segment-specific dynamics inaccessible to conventional approaches. Our findings show that, in the active state, the βarr1 tail preferentially engages the back side of the main body and forms substates in which the middle segment occupies the central crest crevice, suggesting that the released tail can self-engage functionally critical surfaces and influence the balance between tail-only and core-engaged receptor complexes. Together, this work establishes a TAE framework for analyzing large-scale conformational ensembles and advances our understanding of βarr1 activation.

## INTRODUCTION

G protein-coupled receptors (GPCRs) are integral membrane proteins that regulate diverse cellular and physiological processes and represent targets for more than one-third of all FDA-approved drugs (Hauser et al. 2017, Sriram and Insel 2018). Upon ligand binding, GPCRs activate cognate G proteins on the cytoplasmic side to trigger intracellular signaling cascades (Weis and Kobilka 2018). In parallel, many GPCRs engage non-visual β-arrestins (βarr1 and βarr2, collectively βarr), which mediate G protein desensitization and promote clathrin-dependent receptor internalization (Peterson and Luttrell 2017, Gurevich and Gurevich 2019). Beyond these canonical functions, receptor-bound βarr scaffold kinases and signaling partners to initiate distinct G protein-independent pathways (Smith et al. 2018, Ahn et al. 2020). A central mechanism proposed to confer specificity on the βarr-mediated outcomes is the phosphorylation barcode hypothesis, whereby distinct patterns of GPCR phosphorylation are decoded by βarr to favor different βarr conformational states, trafficking responses, and signaling outputs (Nobles et al. 2011). Understanding the mechanisms of βarr activation and pathway selectivity is critical for exploiting their therapeutic potential (Xu and Shao 2022).

Arrestins are composed of two structured domains, the N and C domains, which together form the main body of the protein. The interdomain loops connecting these domains create the central crest region. In the basal state, the C-terminal tail of βarr (βarr tail) engages a groove on the N domain (N-domain groove), resulting in autoinhibition (Han et al. 2001, Asher et al. 2022). Specifically, the middle segment of the βarr tail adopts an extended conformation within this groove (Asher et al. 2022). Upon receptor activation, the phosphorylated receptor tail (Rp tail), enriched in negatively charged phosphate groups on Ser or Thr residues, binds to the positively charged N-domain groove, displacing the βarr tail and thereby promoting and stabilizing the active state of βarr. This displacement exposes clathrin- and adaptin-binding sites in the βarr tail to facilitate receptor endocytosis, while Rp-tail engagement drives activation-associated rearrangements of the βarr main body that contribute to receptor engagement and the assembly of downstream signaling complexes (Xiao et al. 2007). Therefore, mechanistic insights into the functionally relevant conformational ensemble of the βarr tail and its spatial relationship to the βarr main body and the receptor are crucial (Gurevich and Gurevich 2015, Perry-Hauser et al. 2022). Furthermore, distinguishing the impact of the bound Rp tail from that of the βarr tail on the structural rearrangements of the main body is critical for understanding how distinct interactions within the N-domain groove shape the conformational landscape of βarr activation.

However, the positions, conformations, and dynamics of the highly flexible βarr tail relative to the main body in the active state are largely unknown, as the βarr tail was either removed or unresolved in the experimentally determined structures, likely owing to its the dynamic and at least partially unstructured nature (Gurevich et al. 2018, He et al. 2021). Nevertheless, a recent study revealed that when the Rp tail of the atypical chemokine receptor 2 (D6Rpp) binds to βarr2, the middle segment of the βarr2 tail shifts from a β-strand conformation in the N-domain groove to a helical conformation upon engaging the central crest pocket at the interface of a βarr2 dimer. This structural adaptability has led to the middle segment of the βarr tail being referred to as the chameleon motif. Notably, occupation of the central crest pocket by this motif would be expected to prevent receptor core engagement (Maharana et al. 2024). Nevertheless, it remained unclear whether this helical engagement with the crevice represents the only binding mode for the chameleon motif, whether it is specific to D6Rpp, how the rest of the βarr2 tail is positioned relative to the main body, and to what extent analogous interactions are conserved or diverge between βarr2 and βarr1.

While molecular dynamics (MD) simulation is a suitable approach to sample and characterize the conformational ensemble of the 62-residue βarr1 tail, this long tail can move extensively around the main body and form transient secondary structures within a large three-dimensional space. Achieving adequate sampling of this landscape lies beyond the timescales accessible to conventional MD simulations. This limitation can be addressed using enhanced sampling techniques such as temperature replica-exchange molecular dynamics (TREMD), which substantially improve sampling efficiency within practical computational times (Sugita and Okamoto 1999, Rhee and Pande 2003). However, the large and complex datasets produced by TREMD simulations complicate downstream analysis when relying solely on conventional geometric measures.

Recent advances in machine learning have opened new avenues for analyzing high-dimensional molecular simulation data, enabling the extraction of physically meaningful patterns that are often inaccessible through conventional statistical or geometric approaches (Noe et al. 2020). Neural network-based methods, including autoencoders and graph neural networks, have proven effective in learning latent representations that capture collective motions, reaction coordinates, and conformational transitions from complex molecular ensembles (Mardt et al. 2018, Wehmeyer and Noe 2018, Noe et al. 2020). More recently, transformers have emerged as a particularly powerful class of neural network models due to their self-attention mechanism, which allows them to capture long-range dependencies (Vaswani et al. 2017). Building on this capability, we asked whether transformer-based models could learn context-aware relationships among molecular features directly from conformational ensembles.

In this study, we developed a transformer-based autoencoder (TAE) framework to analyze large-scale TREMD data and investigate the conformational organization and activation-linked dynamics of the βarr1 tail. This unsupervised learning approach enables residue-level couplings and modular organization to be inferred directly from simulation data without relying on predefined structural motifs or metrics. We demonstrate that this framework can systematically characterize the heterogeneous conformations of flexible protein regions and identify conformational clusters that are otherwise difficult to resolve using conventional clustering approaches.

## RESULTS

As described in the companion paper, full-length models of βarr1 in both basal and active states were constructed using crystal structures of arrestins as templates, with unresolved regions built through homology and *ab initio* modeling (see Methods in (Ngo et al. 2025)). To enhance conformational sampling, temperature-replica exchange molecular dynamics (TREMD) simulations were then performed for both states (**Table S1**) (Ngo et al. 2025). In the TREMD simulations of the V2Rpp-bound active state of βarr1, the dissociation of the βarr1 tail from the N-domain groove allowed it to explore a vast conformational space, yielding a highly diverse conformational ensemble (Ngo et al. 2025).

### A transformer autoencoder framework for uncovering βarr1 tail modularity and conformational states

To overcome the limitations of conventional approaches in characterizing the βarr1 tail conformations, we developed a transformer-based autoencoder (TAE) framework that analyzes TREMD ensembles in an unsupervised and systematic manner to learn intrinsic residue-level couplings and collective motions of the βarr1 tail.

For each simulation frame, the Cα-Cα distances between each βarr1 tail residue and all other residues (including other tail residues) were computed to capture the internal topology of the protein while remaining invariant to global translation or rotation, i.e., independent of structural superposition. This representation enabled the model to treat each tail residue as an individual token while capturing the collective geometry of the tail within the context of the entire protein (see Methods).

To learn compact representations of βarr1-tail conformations, TAEs were then trained to reconstruct these residue-distance patterns. The encoder was implemented as a transformer with stacked self-attention layers, which transformed residue-wise inputs into contextualized residue representations. A pooling-by-multi-head-attention (PMA) readout then condensed these representations into a small number of slot representations, which were passed through a latent bottleneck to produce compact frame-level embeddings. Using these embeddings as memory, the transformer decoder reconstructed the original residue-distance patterns, and the model was trained by minimizing the reconstruction loss (**Fig. 2**). Model parameters, including the input dimensionality, number of transformer layers, and number of pooling slots in the PMA readout, were systematically tuned to improve reconstruction quality and model stability while keeping model complexity manageable (see Methods).

**Figure 2.**
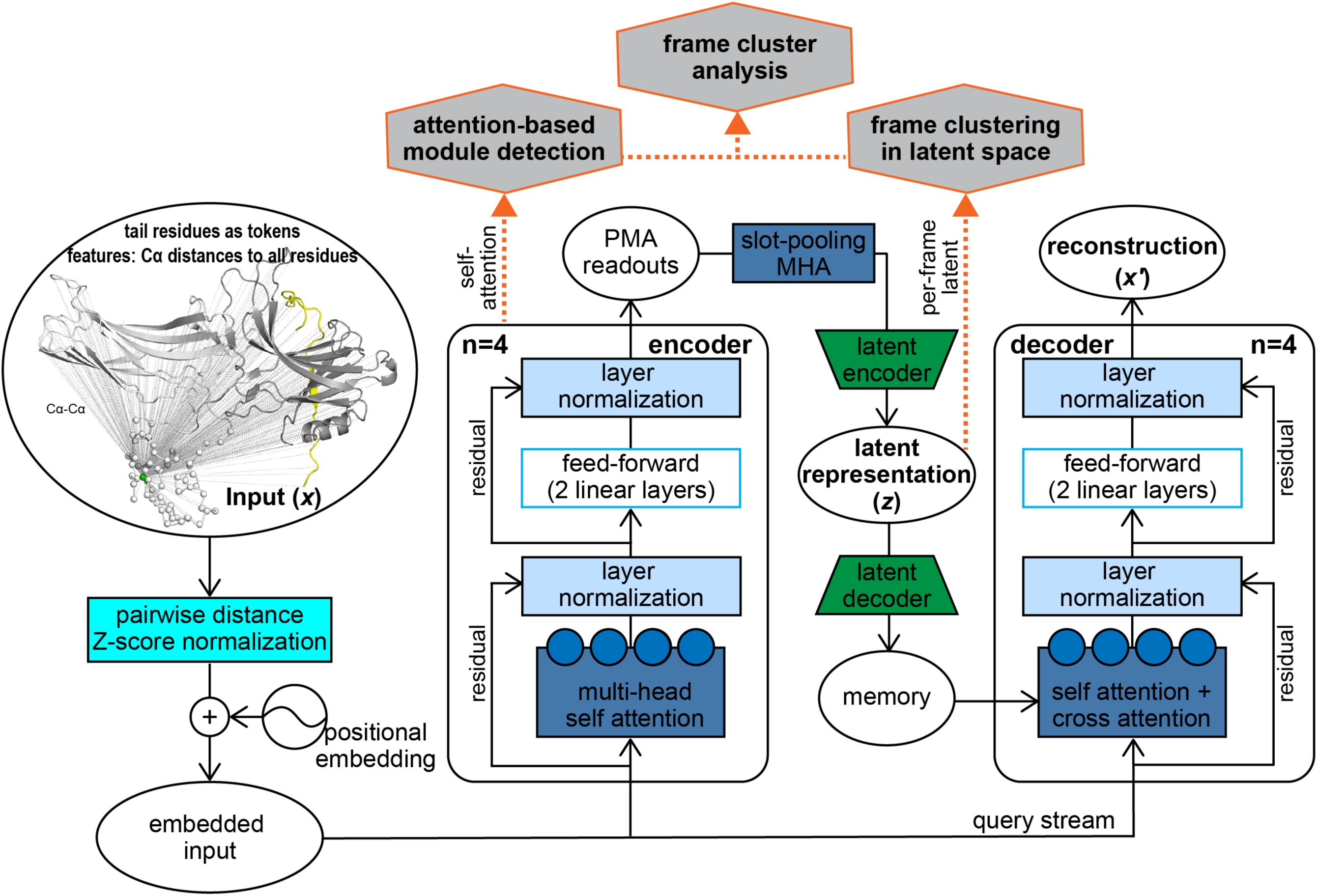
Transformer-based autoencoder architecture for conformational embedding of the βarr tail. Each residue in the βarr tail is represented as a token in the encoder input, and the features associated with each token are its pairwise Cα-Cα distances to all residues. In the molecular graphic depiction, the Cα atoms of the βarr tail residues are shown as spheres; for clarity, the pairwise distances are illustrated for only one tail residue, Ile377, whose Cα atom is colored green. The input x is organized with tail residues as rows and pairwise Cα-Cα distances to all residues as columns. This distance matrix is Z-score standardized and linearly projected into the model embedding space, after which a learnable positional embedding (one vector per token) is added to generate the embedded input (see Methods). In the schematic, rectangles denote computational operations and ellipses denote data tensors. The rounded boxes labeled “encoder” and “decoder” represent individual layers within the corresponding stacks (four layers each). Each encoder layer consists of a multi-head self-attention sublayer and a position-wise feed-forward sublayer (two linear layers with a ReLU activation in between), each followed by residual connection and layer normalization. The filled circles within the attention modules indicate the four attention heads. After the encoder stack, token-level representations are aggregated by pooling-by-multihead-attention (PMA) into three or five slot vectors and then combined into a single summary by a slot-mixing multi-head attention block (MHA, see Methods). A linear latent encoder module compresses this summary to the latent representation (z). A mirrored latent decoder module expands z and broadcasts it across token positions to generate the memory tensor used by the decoder. Within each decoder layer, the embedded input tokens serve as query (Q), key (K), and value (V) tensors for self-attention and as Q for cross-attention, whereas the K and V tensors for cross-attention are supplied by the memory generated from z through latent decoder. The final decoder output is projected back to the input feature dimension to produce the reconstruction x’, which is compared with x during training using a mean squared error objective. Post-training analyses (orange hexagons) are performed with the model in inference mode, with model weights frozen and no additional training. Attention-based module detection: the encoder self-attention maps are averaged over frames, layers, and heads, converted to a distance, and agglomeratively clustered to partition C-tail residues into structural modules (see Methods). Frame clustering in latent space: one latent vector is collected per frame; these vectors are L2-normalized, reduced with UMAP, and clustered with HDBSCAN, using KMeans as a complementary check (see Methods). Frame cluster analysis: the resulting frame clusters are then interpreted in the context of the modules detected for each condition (basal or active; see Results).

After training, the TAE produced two complementary representations of the conformational ensemble. Averaged attention weights across layers and heads yielded interpretable residue-residue attention matrices, which were subjected to hierarchical clustering to delineate contiguous structural modules within the tail, thereby identifying the regions that move cooperatively or relatively independently across the ensemble. In parallel, the compact latent embeddings captured the overall tail conformation of each simulation frame. Clustering these embeddings revealed distinct tail conformations, which were then interpreted in the context of the attention-derived modules for each condition, providing a low-dimensional characterization of the conformational landscape.

### Transformer analysis reveals modular architecture of the basal βarr1 tail consistent with middle-segment anchoring

We first applied the TAE framework to the basal-state simulations, which were collected as described in the companion study as well (Ngo et al. 2025). In the basal autoinhibited state, the middle segment of the βarr1 tail remained largely anchored in the N-domain groove, restricting the tail to a significantly smaller and more tractable conformational space than in the active state. We therefore used the basal ensemble to test whether the TAE analysis could recover distinct and intuitively interpretable βarr1-tail conformations before applying the same framework to the more heterogeneous active-state ensemble.

After training the TAE on the basal-state ensemble, the derived residue-residue attention matrix was subjected to hierarchical agglomerative clustering to identify the modular organization of the tail (see Methods). High attention between two residues indicates that the model treats their states as jointly informative for reconstructing the tail geometry, reflecting an underlying structural or dynamical coupling. Clustering of the averaged attention patterns therefore reveals cohesive groups of residues that behave as modular units (Roshan et al. 2021) (see Discussion). To avoid bias from any specific training–validation split, 10 replicas were randomly selected from replicas 35–46 (306.9–312.4 K) for training, with the remaining 2 replicas used for validation. This procedure was repeated for all possible 10–2 training–validation splits among these 12 replicas, yielding a total of 66 independently trained models. Clustering the attention matrix from each model separately, we found that, although some local attention patterns varied among individual models, the overall module boundaries defined by clustering remained largely consistent across all 66 models. Ensemble-variability analysis further showed that model-to-model variability was concentrated within dominant mean-attention regions rather than scattered across unrelated regions (**Fig. S2**), supporting the robustness of the attention-derived module-level organization. To suppress local noise and extract the consensus modular structure, we averaged the attention matrices across all 66 models (**Fig. 3A**).

**Figure 3.**
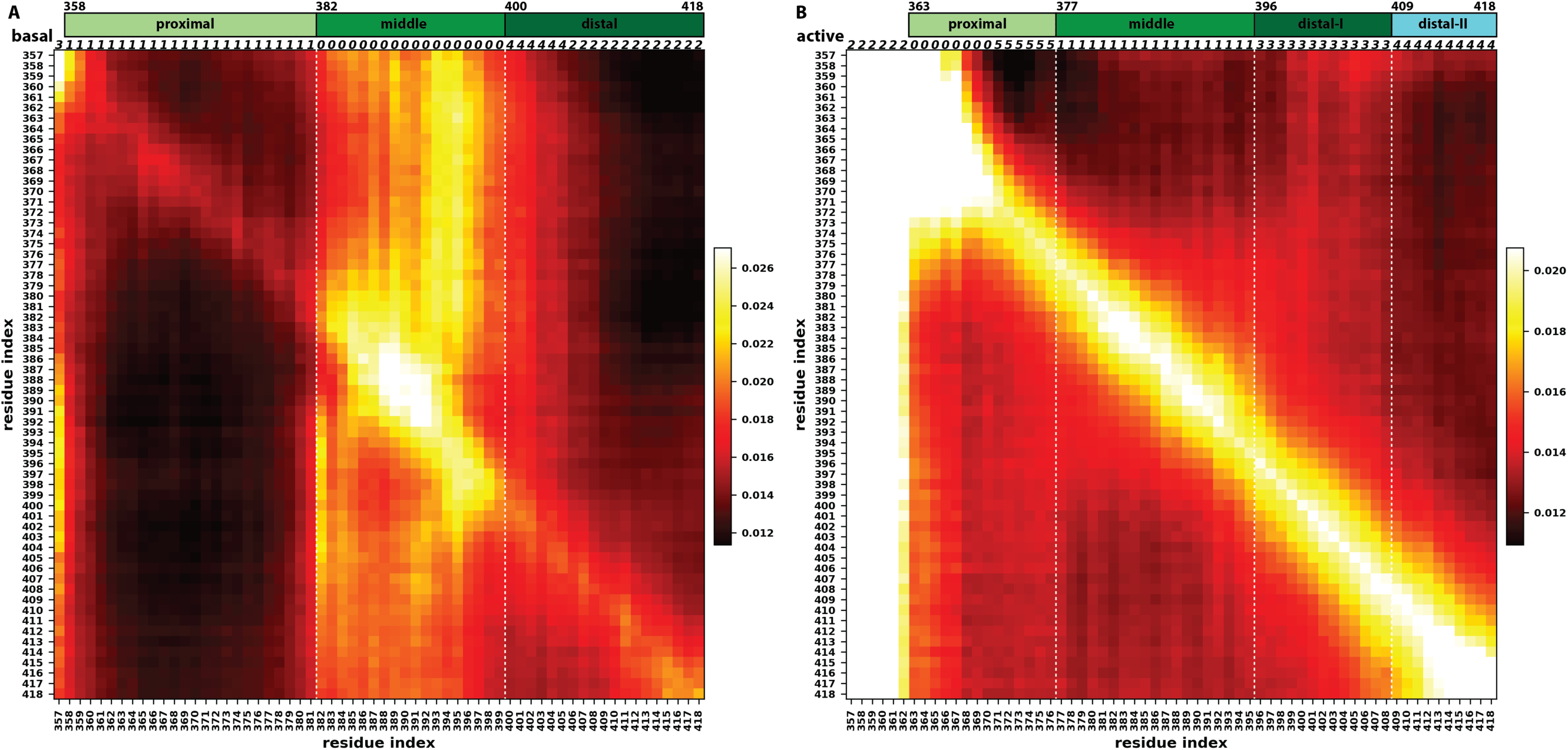
Averaged attention matrices reveal altered βarr1 module organization in basal and active ensembles. Panel A shows the basal-state ensemble-averaged attention matrix as a heatmap of residue–residue attention values averaged across the independently trained basal-state TAE models (see Methods). Both rows and columns correspond to tail residues 357–418. Rows represent query residues and columns represent key residues; thus, each residue on the y-axis attends to residues along the x-axis. The numbers above the top edge of the matrix denote the cluster assignments obtained by hierarchical agglomerative clustering of the averaged attention patterns (see Methods), and the colored bars indicate the three modules (proximal, middle, distal) defined by this clustering. Residue 357 was excluded from the proximal segment because of its hinge role. To maintain a minimum segment length of at least 10 residues, clusters labelled 4 and 2 in the clustering solution were merged into the distal segment. Panel B shows the active-state ensemble-averaged attention matrix constructed using the same model-averaging and clustering pipeline applied to the basal ensemble. In contrast to the basal state, release of the tail from the N-domain groove in the active state redistributes attention: attention outside the hinge region becomes weaker and more diffuse, whereas hinge residues dominate the attention map. Despite this weaker non-hinge attention pattern, applying the same clustering protocol used for the basal state resolved four distinct modules (proximal, middle, distal-I, and distal-II) in the active-state attention matrix, indicated by the colored bars. Residues 357-362 was excluded from the proximal segment because of their hinge role. To maintain a minimum segment length of at least 10 residues, clusters labelled 0 and 5 in the clustering solution were merged into the proximal segment.

Clustering this averaged matrix identified three contiguous modules, closely corresponding to our previous segmentation based on root-mean-square fluctuation analysis (Asher et al. 2022): a proximal segment (residues 358–381) near the hinge, a middle segment (residues 382–399), and a distal segment (residue 400-418) (**Fig. 3A**). This modularization defined by TAE attention encodes a key structural feature, namely anchoring of the middle segment within the N-domain groove. Interestingly, residues 382–384 and 398–399, which were previously grouped with the proximal segment, are now clustered with the middle segment, indicating that the transformer recognized their coupling to the groove-anchoring region despite their flexibility and location outside the groove. Having characterized the modular organization through attention analysis, we next examined the conformational landscape using the latent-space embeddings.

### Latent-space clustering reveals modular flexibility of the βarr1 tail

Because the autoencoder compresses each frame through a low-dimensional bottleneck, it captures the most informative aspects of tail motion in a compact representation amenable to clustering and ensemble-level comparison. These low-dimensional latent embeddings therefore provide a quantitative characterization of the conformational landscape of the βarr1 tail. For the basal-state analysis, frames from the temperature replicas closest to 310 K (replicas 37–46) were passed through the TAE model that achieved the lowest tail reconstruction loss to obtain per-frame latent embeddings, which captured the essential degrees of freedom of tail motion near the physiological temperature.

The latent embeddings were then subjected to a combined Uniform Manifold Approximation and Projection (UMAP) and Hierarchical Density-Based Spatial Clustering of Applications with Noise (HDBSCAN) analysis, in which an extensive grid search over UMAP dimensionality-reduction and HDBSCAN clustering parameters was performed to identify the configuration that maximized the Density-Based Clustering Validation (DBCV) score while constraining the noise fraction to remain low (see Methods). The best-performing parameters were subsequently applied to both the original and PCA-preprocessed embeddings to determine whether denoising the latent space with PCA improved clustering performance (see Methods). The best HDBSCAN clustering using the optimal UMAP embedding produced a silhouette score of 0.589 with all samples included, and a corresponding DBCV score of 0.636, supporting robust density-based structure in the embedding space. After excluding noise points (13.3% of samples) and recalculating the silhouette score on the clustered subset, the modified silhouette score increased to 0.805, indicating strong separation among core clusters. This analysis identified 280 distinct conformational clusters of the basal-state βarr1 tail.

To interpret the latent-space clusters in the context of the attention-derived modular organization, we calculated average pairwise RMSDs of frames within each cluster for the top 40 largest clusters, which account for 30% of the entire basal dataset, evaluating the entire βarr1 tail as well as the proximal, middle, and distal segments separately (**Fig. 4A**). To distinguish internal tail or segment conformational variability from changes in orientation relative to the main body, pairwise RMSDs were computed using two alignment schemes: (i) aligning the structured regions of the main body between two frames before computing the RMSD of the tail or its segments (main-body–tail scheme), and (ii) aligning only the residues included in the RMSD calculation (tail–tail scheme). When plotting RMSDs from the main-body–tail scheme against those from the tail–tail scheme for each region, the values for the entire tail, the proximal and the middle segments were largely linearly correlated (**Fig. 4A**). As expected for the basal state, the anchored middle segment exhibited markedly lower RMSDs across all top clusters in both schemes. In contrast, the linear relationship between the two alignment approaches was noticeably weaker for the more flexible distal segment (**Fig. 4A**), suggesting that their variability arises from changes in both local tail conformation and the orientation of the tail relative to the main body. Thus, our TAE framework captured both sources of variability by jointly encoding intra-tail geometry and tail–body context.

**Figure 4.**
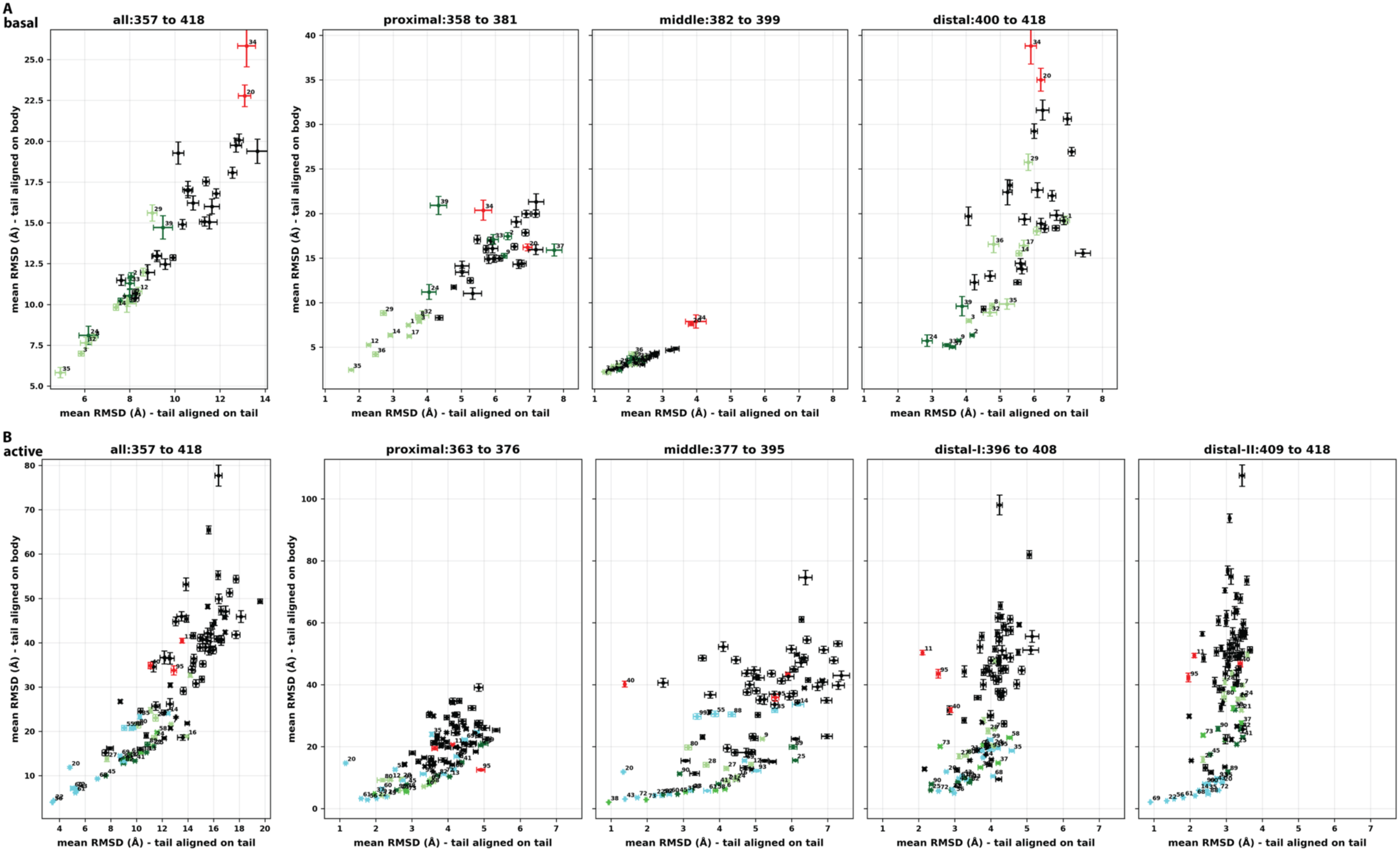
Alignment-scheme comparison reveals segmental decoupling in basal and active βarr1 ensembles. Panel A shows, for the basal ensemble, scatter plots of average intra-cluster pairwise RMSDs for HDBSCAN clusters calculated under two alignment schemes: main-body–tail alignment (y-axis), in which frames were aligned on the structured βarr1 main body before RMSD calculation, and tail–tail alignment (x-axis), in which alignment was performed using only the tail residues included in the RMSD calculation. The left plot corresponds to the entire tail (residues 357–418), and the right plots correspond to the proximal, middle, and distal segments defined by the basal attention-based clustering (Fig. 3). Clusters near the diagonal exhibit variability dominated by internal conformational differences, whereas points positioned substantially above the diagonal indicate additional variability arising from changes in tail orientation relative to the main body. Clusters with the lowest main-body–tail RMSDs in the distal segment (b2, b9, and b24; dark green) display comparatively large RMSDs in the proximal segment. Conversely, among the clusters with the lowest main-body–tail RMSDs in the proximal segment (light green), b12, b14, and b17 display comparatively large RMSDs in the distal segment (see Fig. 5). Panel B presents the same analysis for the active (V2Rpp-bound) ensemble using segment definitions derived from active-state attention clustering. A similar pattern of segmental decoupling is observed in the active state (see **Fig. S5**), but some clusters also exhibit large main-body–tail RMSDs together with low tail–tail RMSDs for specific segments (red clusters, see **Fig. S6**), a feature not apparent in the basal state. All RMSD calculations were performed using the same trajectory frames used for latent-space clustering, and the top 40 and top 100 clusters by population are shown for the basal and active states, respectively.

Interestingly, clusters that are relatively compact in the proximal segment (i.e., exhibiting lower RMSDs) are not necessarily compact in the distal segment, and vice versa. For examples, the clusters with the lowest main-body–tail RMSDs in the distal segment (clusters b2, b9, and b24; “b” denotes basal) all display comparatively large RMSDs in the proximal segment, whereas the clusters with the lowest main-body–tail RMSDs in the proximal segment (clusters b12, b14, and b17) all show comparatively large RMSDs in the distal segment (**Fig. 5A,B**).

**Figure 5.**
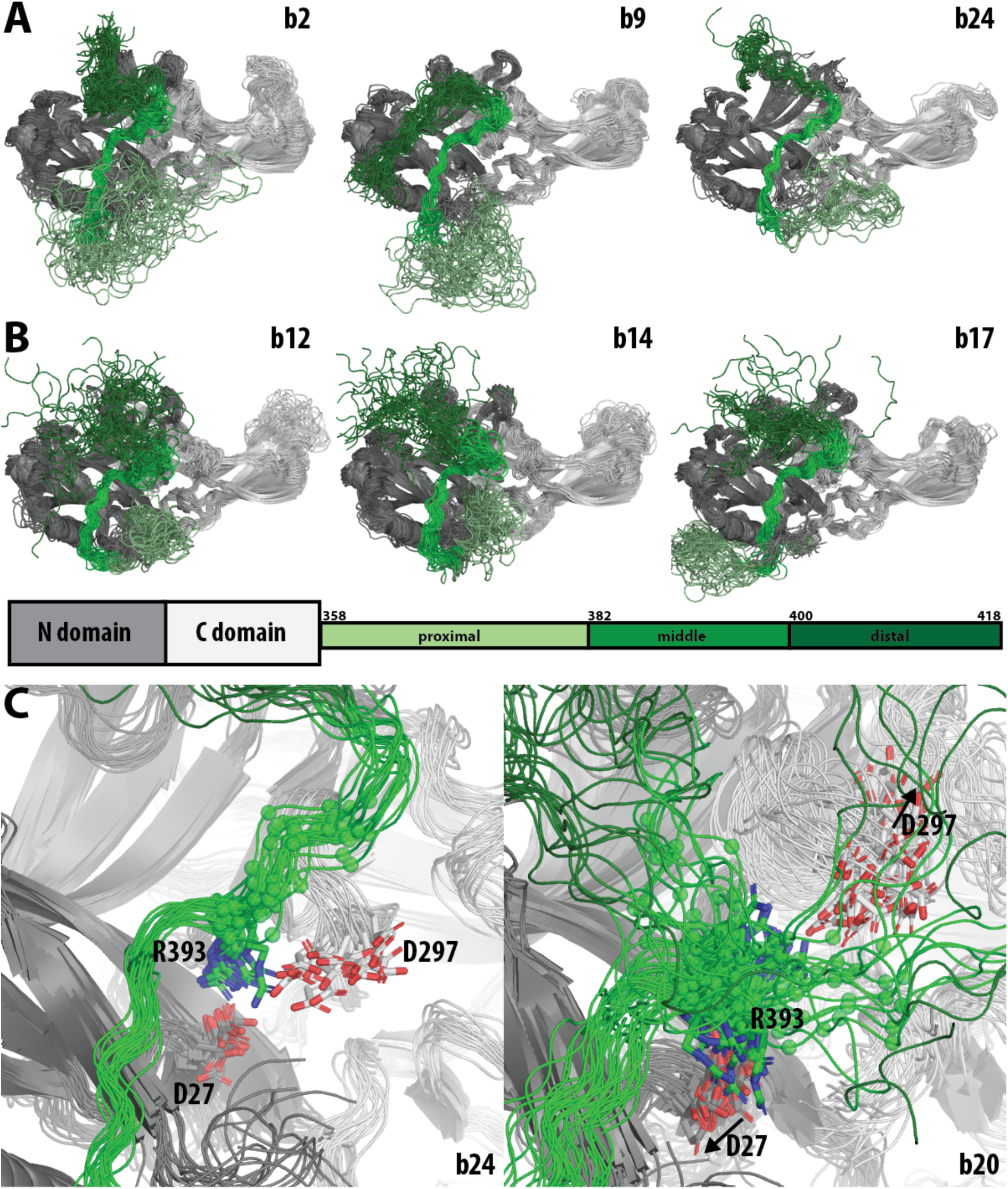
Selected basal-state cluster conformations illustrating segmental decoupling of tail organization. Panel A shows that the clusters with the lowest main-body–tail RMSDs in the distal segment (clusters b2, b9, and b24, see Fig. 4A) nevertheless exhibit high mobility in the proximal segment. In these clusters, the distal segment occupies the extended groove, although in distinct configurations. Panel B shows the converse pattern: in clusters with low main-body–tail RMSDs in the proximal segment (clusters b12, b14, and b17), the proximal segment is tightly clustered, whereas the distal segment is dispersed. The color assignment of each domain and segment is indicated in the schematic below panel B. Panel C shows that cluster b24, representative most clusters, retains tight engagement of the middle segment with the N-domain groove and neighboring regions, including the polar core. In contrast, cluster b20 displays both elevated main-body–tail RMSDs and increased tail–tail RMSDs, consistent with disengagement of the C-terminal portion of the middle segment from the N domain. Notably, R393 is dissociated from its ionic interactions with D27 and D297 in the polar core.

Examination of representative frames of clusters b2, b9, and b24 revealed that, in these clusters, the distal segment either partially or fully occupies the extended groove on the N domain with various positionings (**Fig. 5A**), a configuration reported to be functionally relevant (Asher et al. 2022) but showing no consistent correspondence with any defined conformational state of the proximal segment. These observations suggest a decoupling between the proximal and distal regions, consistent with a modular organization of the βarr1 tail. This interpretation aligns with the attention analysis, which showed minimal cross-attention between proximal and distal residues, indicating weak cooperative coupling between them (**Fig. 3A**).

The cluster with the largest RMSDs in the middle segment (cluster b20) reveals an additional mechanistic feature. In this cluster, the C-terminal portion of the middle segment (residues 393–396), in which R393 forms ionic interactions with D26 and D297 of the so-called “polar core” (Asher et al. 2022), disengages from its interaction with the N domain (**Fig. 5C**), representing a partially engaged intermediate state. Such disengagement was previously observed in our conventional MD simulations of the basal state only with the three-element interaction (3-EI) mutations (I386A/V387A/F388A), which demonstrated that disrupted hydrophobic 3-EI between the middle segment and the N-domain helix allosterically perturbed the association between R393 and the polar core (Asher et al. 2022). This finding suggests that the engagement with the polar core may occur as a mechanistically distinct, though allosterically coupled, event from the engagement with the N-domain groove. Importantly, the emergence of this state in the present TREMD simulations of the wild-type βarr1, without mutations, underscores the ability of replica-exchange sampling to more comprehensively explore the conformational landscape of the βarr1 tail and to uncover transient intermediate states that are rarely sampled in conventional MD.

These subtle yet functionally meaningful features—including local conformational changes like partial disengagement as well as segment-level patterns like proximal-distal decoupling—would be difficult to detect using conventional analyses without predefined metrics, structural boundaries, or prior expectations about where to focus attention. In contrast, the TAE framework resolves these features in an unsupervised manner by capturing both the internal conformational variability of the βarr1 tail and its placement relative to the main body, thereby revealing segment-specific behavior and subtle mechanistic states. Having characterized the structural and dynamical features of the tail in the basal state, we next examined how these properties change upon receptor activation.

### The unrestrained active βarr1 tail exhibits expanded conformational diversity and an altered modular architecture

To determine how the organization and dynamics of the βarr1 tail are reshaped in the active state, we applied the same TAE-based analysis used for the basal state, consisting of attention-derived module identification followed by latent-space clustering of TAE embeddings. We sought to address three questions: (i) whether the tail’s basal-state modules are preserved or reorganize upon release from the N-domain groove; (ii) whether the tail establishes new interactions with the main body that serve as critical anchoring points to constrain tail motion; and (iii) whether functionally relevant tail conformational states occupy only a small fraction of the conformational space sampled in our simulations.

Because the active-state ensemble exhibited substantially more conformational diversity than the basal state and the corresponding dataset was significantly larger, the TAE architecture and model parameters were re-optimized for the active-state data (see Methods). As for the basal state, we trained 66 independent active-state TAE models using all possible 10–2 training-validation splits of replicas 35–46 with the optimized parameters. We first examined the attention patterns they collectively learned.

The model-averaged attention matrix for the V2Rpp-bound active state revealed a markedly different attention pattern from the basal state, resulting in a distinct module division. Specifically, because the βarr1 tail was no longer anchored in the N-domain groove and instead sampled a much broader conformational space surrounding the main body, the distribution of residue-to-residue attention values shifted substantially (**Fig. 3B**). Compared with the basal state, attention from most tail residues to other tail positions was generally diminished, whereas attention toward the hinge residues at the beginning of the tail, which connect the main-body and tail, became highly concentrated. This pattern is expected in the active state, where the hinge region acts as a dominant anchor for orienting the tail relative to the main body, while the remainder of the tail explores a more dispersed set of conformations. Nevertheless, the residual attention structure, although more concentrated near the diagonal and thus indicative of more localized attention, remained sufficient to resolve the tail into segments, as evidenced by the faint red, roughly square patches in **Fig. 3B** that denote nontrivial attention outside the strongest yellow regions. Applying the same hierarchical agglomerative clustering protocol used for the basal state, we found that, despite some resemblance to the basal state, the resolved module layout in the active state showed notable differences, with boundaries shifted and the distal region further subdivided. Specifically, this reorganization resulted in four segments: a proximal segment (residues 363–376, excluding the first six hinge residues 357–362), a middle segment (residues 377–395), and two distal segments, distal-I (residues 396–408) and distal-II (residues 409–418) (**Fig. 3B**). To assess whether the strong attention focused on the hinge region (residues 357–362) biased the attention structure, we retrained the models with the hinge residues excluded from the token set and obtained a similar module organization (**Fig. S3**). We therefore used the models that included the hinge region, as they retain more information, for the subsequent analysis.

### Active-state clustering reveals segmental decoupling and segment-specific conformations

We next performed UMAP–HDBSCAN clustering of the active-state latent embeddings using the same pipeline and grid-search procedure as for the basal state, yielding an optimized parameter set specific to the active-state ensemble (see Methods). The best HDBSCAN clustering of the optimal UMAP embedding produced a silhouette score of 0.606 and a DBCV score of 0.627. After excluding noise points (12.3% of the total samples), the modified silhouette score increased to 0.819. This analysis identified 738 distinct clusters. Additional analysis of the active-state HDBSCAN parameter path and cluster-size distribution supported that this high cluster count was not dominated by parameter-driven over-fragmentation, but instead reflected the intrinsic conformational heterogeneity of the active ensemble (see **Supplemental Results**; **Table S2**; **Fig. S4**). Together, these results indicate a marked increase in conformational heterogeneity relative to the basal state, reflecting greater diversity in both intra-tail conformations and tail–main-body orientations.

Using the active-state module definitions as the interpretive framework for the latent-space clusters, we plotted RMSDs from the main-body–tail scheme against those from the tail–tail scheme for each region, for the top 100 largest clusters, which account for 30% of the entire active dataset (**Fig. 4B**). In contrast to the same analysis for the basal state, only the full tail exhibited a roughly linear correlation trend, while individual segments showed markedly weakened relationships between the two alignment schemes, especially for the middle and two distal segments. This indicates that segment-level variability in the active state arises from combined changes in internal conformation and orientation relative to the main body, whose interplay is resolved by our TAE framework.

As observed in the basal state, some clusters exhibit consistently low tail–tail and main-body–tail RMSDs across all segments, whereas others display clear decoupling between segments. For example, clusters a7 (“a” denotes active) shows well-defined proximal and middle region paired with a considerably more dispersed distal-II region, whereas cluster a9 shows a relatively well-defined proximal region but is markedly more variable for all other segments. By comparison, cluster a14 and a35 exhibit low distal-II main-body–tail RMSDs, yet their middle segments are relatively dynamic (**Figs. 4B** and **S5**).

Interestingly, several clusters show relatively large main-body–tail RMSDs, yet display relatively low tail–tail RMSDs within specific segments. This pattern indicates that those segments have relatively defined conformations even when being relatively dissociated from the main body. For example, cluster a20 has low tail–tail RMSDs for the proximal and middle segments (1.2 and 1.3 Å, respectively) but relatively large main-body–tail RMSDs for the same segments (14.7 and 11.8 Å, respectively). Visual examination showed that, although the tail is generally located beneath the N domain in this cluster, it does not adopt a tight conformation relative to the main body (**Fig. S6A**). However, when the frames of this cluster are superimposed using the proximal and middle segments, a well-defined structure emerges: residues 388-395 at the end of the middle segment form two helical turns, the rest of the segment (residues 377-387) pack against these two turns, and the proximal segment (residues 363–376) encircles the other side of the helical turns (**Fig. S6B**). In comparison, cluster a40 shares a similar conformation of the middle segment (residues 377–395) with a20, but not of the proximal segment (**Fig. S6D**). This cluster has a tail–tail RMSD for the middle segment of 1.4 Å despite main-body–tail RMSDs of more than 32 Å for the middle, distal-I, and distal-II segments, consistent with broad spatial sampling of the tail without close association with the main body (**Fig. S6C**). These results show that because the middle segment is restrained in the basal state within the N-domain groove in a relatively extended conformation, some of its conformational behaviors become accessible only in the active ensemble. In contrast to clusters a20 and a40, clusters a11 and a95 exhibit defined conformations in the distal-I and distal-II segments but are unstructured in the proximal and middle segments, with neither adopting a clear orientation relative to the main body (**Fig. S6E–H**).

These results further demonstrate the superiority of the TAE models over conventional RMSD-based analyses in recovering structurally interpretable conformations without prior knowledge of which structural features to monitor, while the distinct patterns of segmental structural organization observed in clusters a20 and a40 versus a11 and a95 also align with the module division in the attention analysis.

### Unsupervised TAE clustering recapitulates manually identified interaction patterns and reveals additional tail–main-body variation

To validate whether the TAE-derived clusters recapitulate the interaction patterns identified through manually guided clustering (see the first Results section), we measured for each cluster the distances of the same residue pairs identified by the pairwise contact analysis (see above): Thr374–Arg307 (proximal segment to back side), Asn375–Met255 (proximal segment to front side), and Ala392–Phe244 (middle segment to central crest crevice). Consistent with the manually guided analysis, the TAE clustering revealed a strong bias for the tail to interact with the main body along the back side. Specifically, among the top 100 clusters, 28 had the proximal segment not located at the bottom (defined as having at least one of the Thr374–Arg307 and Asn375–Met255 distances below 22 Å), and 23 were closer to the back side than to the front side (**Fig. S7A**). Visual inspection confirmed that in these clusters, the tail extends upward along the back side of the main body.

Among these 23 clusters, 11 showed the middle segment approaching or entering the central crest crevice (Ala392–Phe244 distance <17 Å; **Fig. S7B**). Structural examination revealed that these 11 clusters differ in how the tail engages the main body along the back side and, in part as a consequence, in how the middle segment is associated with different orientations of the finger loop and middle loop of the N domain and the C-loop of the C domain, which together form and dynamically shape the crevice. **Fig. 6** and its legend provide a comprehensive description and comparison of these clusters. Here, we focus on selected clusters that exhibit tighter tail–main-body engagement.

**Figure 6.**
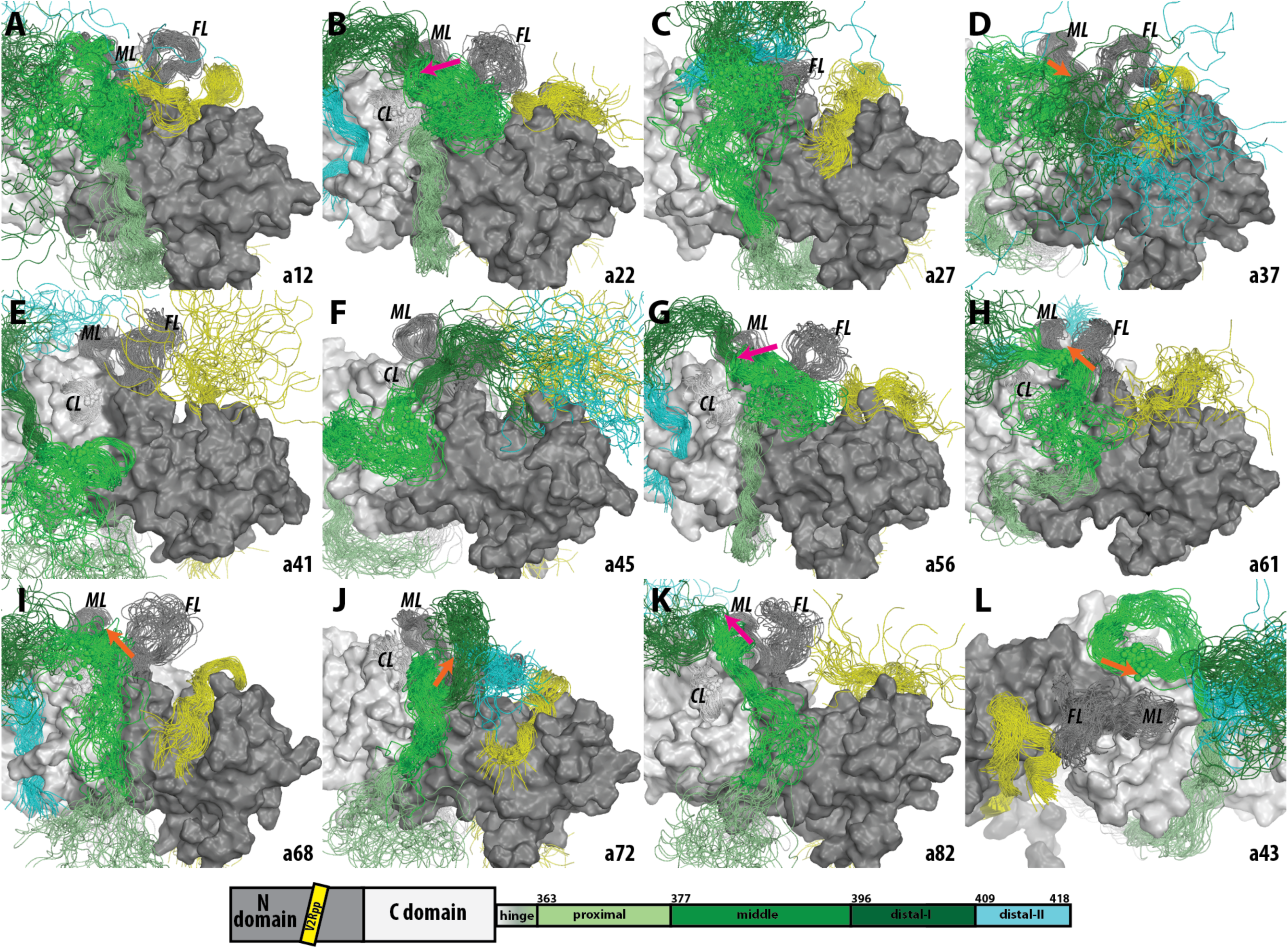
Clusters among the top 100 HDBSCAN clusters with the middle segment in the vicinity of the central crest crevice. 12 clusters were identified using a distance criterion between Ala392 and Phe244, defined as a Cα-Cα distance below 17 Å (see **Fig. S7**). Of these, 11 approach the crevice from the back side (panels A-K), whereas one approaches from the front side (panel L). Panels A-J are shown from the same back-side view of the main body, whereas panel K is shown from the front-side view. The Cα atoms of Ala392 and Phe244 are shown as spheres. The central crest crevice is formed by the finger loop (FL), middle loop (ML), and C-loop (CL). Cluster IDs are shown at the bottom right of each panel. The color assignment of each domain and segment is indicated in the schematic at the bottom. Magenta and orange arrows indicate the orientation of the middle segment within the central crest crevice; clusters without arrows do not show middle-segment insertion into the crevice. Magenta indicates conformations in which residues 388–395 form two helical turns and orange indicates those in which they do not. In cluster a41 (E), the middle segment approaches the crevice without inserting into it, whereas in cluster a12 (A) it generally lies over the C-loop, with the finger loop twisted such that the bound V2Rpp peptide partially occupies the crevice. In cluster a27 (C), while the middle segment is in the vicinity of the crevice, it is the distal-II segment that inserts into the crevice. Clusters a68 (I) and a72 (J) both show upward engagement of the proximal and middle segments along the interface between the N and C domains, but differ in the distal-II segment, which engages the back groove in a68 and partially occupies the crevice while contacting the finger loop in a72. Clusters a22 (B) and a56 (G) both show back-to-front entry of the middle segment into the crevice, with residues 388–395 forming helical turns perpendicular to the finger loop and the distal-II segment engaging the back groove; a56 closely resembles a22 in tail conformation but is associated with a distinct main-body conformation (**Fig. S8**). Unlike a22, a56, and a68, which occupy the back groove with the distal-II segment, a37 (D) and a45 (F) engage this groove with the proximal segment, while their middle segments approach the central crest crevice from different directions, resulting in distinct special distributions of the distal segments. Cluster a82 (K) shows back-to-front entry of the middle segment into the central crest crevice, with residues 388–395 forming two helical turns parallel to the finger loop and aligned with the corresponding region of the βarr2 chameleon motif. Cluster a61 (H) shows some similarity to a82 in the location and orientation of the middle segment, but lacks these helical turns; in contrast to the relatively disordered distal-II segment in a82, the distal-II segment in a61 tightly engages the groove on the front side of the C domain (“front groove”). Cluster a43 (L) is the only top-100 cluster that approaches the central crest crevice from the front side of the C domain, with the middle segment passing behind the C-loop and entering the crevice in a back-to-front orientation without forming helical turns.

Clusters a68 and a72 show some similarity in how their proximal and middle segments engage the main body, extending upward toward the central crest crevice along the interface between the N and C domains, with the last four to five residues of the middle segment entering the crevice. However, their distal-I and distal-II segments adopt opposite orientations. As a result, in a68 the distal-II segment engages the groove on the back side of the C domain (hereafter termed the “back groove”), whereas in a72 it partially occupies the crevice and interacts with the finger loop (**Fig. 6I,J**).

In clusters a22 and a56, the middle segment enters the central crest crevice in a back-to-front orientation, with the helical turns formed by residues 388–395 oriented largely perpendicular to the finger loop (**Fig. 6B,G**). These clusters also resemble a68 in their overall orientation and close association with the main body, including engagement of the distal-II segment with the back groove, although their proximal tails are positioned quite differently from that in a68. Notably, a22 and a56 have very similar tail conformations; however, superposition and visual inspection suggest that they are associated with distinct global conformations of the main body. Analysis with the protein interaction analyzer (PIA) (Stolzenberg et al. 2016, Ngo et al. 2025) indicated that the most apparent differences occur near the arrestin switch I (ASwI) and α-helix I within the three-element interaction region associated with V2Rpp binding (**Fig. S8**). Because these two clusters show the most extensive tail–main-body interactions, the divergence in the main body produced two separate clusters even when the tail adopted nearly the same bound configuration. In the unsorted TREMD trajectory from which these clusters arise, a56 occurs immediately after a22, raising the possibility that the transition from a22 to a56 reflects a change in main-body conformation induced by anchoring of the βarr1 tail in this tightly associated configuration. Thus, the separation of a22 and a56 into two clusters suggests that the TAE-based clustering captures not only differences in the tail conformation, but also the main-body variation that may result from close βarr1 tail binding.

### Cluster a82 captures a shortened βarr2-like chameleon-motif helix in the central crest crevice

For cluster a82, the middle segment enters the central crest crevice in a back-to-front orientation (**Figs. 6K** and **7A**), running parallel to the finger loop. Remarkably, this configuration from our V2Rpp-bound βarr1 simulations matches that of the chameleon motif in the crevice of the D6Rpp-bound βarr2 (**Fig. 7B**). However, in a82, residues 388–395 of the βarr1 chameleon motif form only two helical turns, whereas the βarr2 chameleon motif forms four and a half helical turns in the crevice. Nevertheless, superposition shows that these two turns align well with the corresponding region near the end of the four-and-a-half-turn helix in the βarr2 structure, adopting the same orientation and occupying the same region of the central crest crevice (**Fig. 7**), consistent with a common mode of crevice engagement.

**Figure 7.**
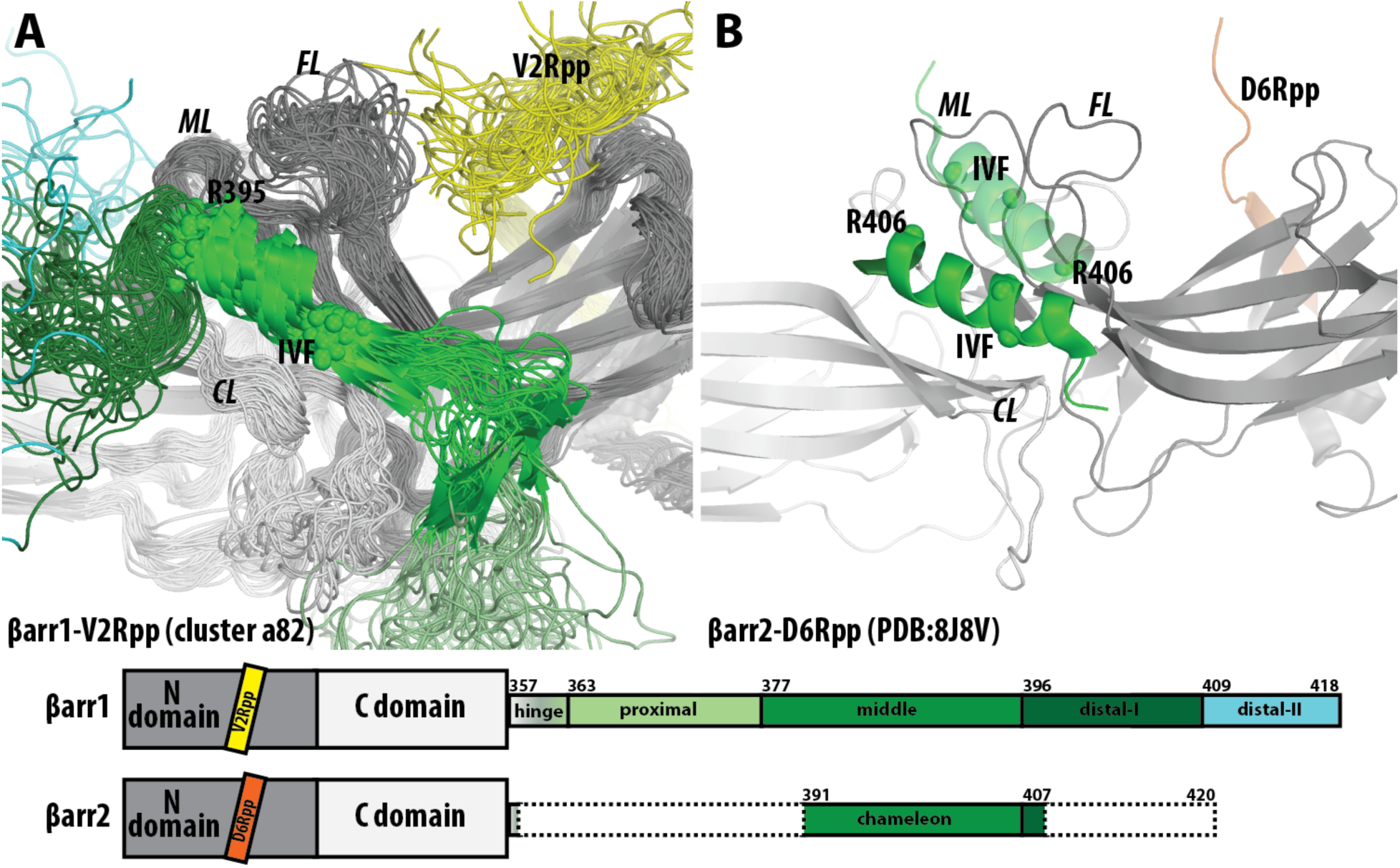
The βarr1 tail in cluster a82 recapitulates engagement of the central crest crevice by the chameleon motif in D6Rpp-bound βarr2. Panel A shows a zoomed-in view of βarr1-V2Rpp cluster a82, in which the middle segment enters the central crest crevice in a back-to-front orientation, parallel to the finger loop (FL). Residues 388–395, within the chameleon motif region of βarr1 middle segment, form a short two-turn helix in the crevice; the preceding consecutive hydrophobic residues IVF (residues 386–388) adopt a β-strand conformation, as in the N-domain groove in the basal state. Residue 386-388 and R395 are marked by Cα spheres. Panel B shows the aligned D6Rpp-bound βarr2 structure (PDB: 8J8V), in which the chameleon motif forms a four-and-a-half-turn helix in the same crevice; the Cα atoms of the corresponding IVF residues and R406 are shown as well. Structure 8J8V is a βarr2 dimer in which the chameleon-motif helix forms part of the dimer interface through interaction with the corresponding helix of the other monomer. For simplicity, only the chameleon motif of the other monomer is shown transparently, with the Cα atoms of its IVF residues and R406 shown as spheres. Comparison of the two panels shows that the two helical turns in cluster a82 align with the corresponding region near the end of the longer βarr2 helix, occupying the same region of the central crest crevice and adopting the same orientation, consistent with a shared mode of crevice engagement. The color assignments of each domain and segment for both βarr1 and βarr2 are indicated in the schematic at the bottom.

The longer helical extent in βarr2 is consistent with the fact that the βarr2 cryo-EM structure was resolved as a dimer, in which the chameleon motif contributes extensively to the dimer interface, thereby shielding its hydrophobic residues from solvent. In particular, the three consecutive hydrophobic residues IVF (βarr1 residues 386–388), which mediate the conserved three-element interaction in the basal state, are helical in the βarr2 structure but not in our βarr1 simulations, suggesting that this intermolecular packing is at least partly responsible for stabilizing the longer helical conformation. Consistent with this interpretation, **Fig. S6**, as described above, shows that residues 388–395 can form a helix even in the absence of association with the main body in clusters a20 and a40, whereas the neighboring residues do not. This observation supports the idea that the chameleon motif in βarr1 has an intrinsic tendency to form only a shorter helical segment rather than the more extended helix observed in dimeric βarr2. This interpretation, however, does not preclude βarr1 oligomerization, which has been linked to cellular localization (Milano et al. 2006), but rather emphasizes the specific dimer-interface context of the βarr2 structure.

Cluster a43 is the only cluster among the top 100 clusters that was able to approach the crevice from the front side of the C domain. Interestingly, its middle segment passes behind the C-loop and enters the crevice in a back-to-front orientation, but without forming any helical conformation. This observation further supports the back-to-front orientation for the middle segment to engage with the crevice; however, in a43, achieving this orientation requires the tail to remain relatively extended through the end of the middle segment, making formation of helical turns by residues 388–395 unlikely while retaining crevice engagement (**Fig. 6L**).

Together, these findings show that the central crest region can be approached through multiple routes, with approach from the back side being strongly favored, while the crevice itself accommodates multiple modes of tail engagement. Thus, the unsupervised TAE framework not only recovered the key interaction patterns that previously required manually guided clustering to resolve, including preferential back-side positioning and crevice occupation, but also resolved the underlying conformational heterogeneity within these binding modes in a way that would be difficult to achieve systematically using conventional approaches.

### Middle-segment anchoring near the central crest crevice supports sustained βarr tail–main-body engagement

These findings raise the possibility that the distinct βarr-tail–main-body engagement patterns identified above may represent states along a conformational progression. A formal kinetic analysis of the present TREMD data, however, is not feasible. In the unsorted replica trajectories, each continuous coordinate trajectory undergoes temperature exchanges and therefore does not represent dynamics at a single thermodynamic state. As a result, transition counts from these trajectories cannot be interpreted as unbiased kinetic transitions for conventional Markov state model (MSM) construction (Bowman et al. 2014). Although the transition-based reweighting analysis method (TRAM) can combine transition information collected at different temperatures (Wu et al. 2016), it still requires physically meaningful within-temperature transitions. Here, the available within-temperature segments are too short for reliable lag-time sampling, and concatenating them would introduce nonphysical boundary jumps, thereby precluding TRAM-based kinetic analysis.

Nevertheless, the temperature-unsorted TREMD trajectories, before frames were reassigned to temperature-specific ensembles, can still be inspected to assess how distinct βarr-tail–main-body engagement patterns appear and rearrange during enhanced sampling. We therefore used them to trace, only qualitatively, the apparent progression of major structural events. As expected, our examination showed that initial back- or front-side association of the proximal segment is followed by approach of the middle segment toward the central crest region. This ordering was observed in all representative unsorted trajectories we examined that include the key clusters mentioned above: utraj-29 (including a43; movie S1), utraj-37 (including a61; movie S2), utraj-70 (including a72; movie S3), utraj-100 (including a22 and a56; movie S4), utraj-129 (including a68; movie S5), and utraj-190 (including a82; movie S6).

Interestingly, however, this examination reveals two phenomena that were not evident from the clustering analysis. First, once the middle segment begins to associate with the central crest crevice in a back-to-front orientation, it does not, within the timescale sampled in our simulations, dissociate from the vicinity of the crevice in any of these six trajectories, including utraj-29, in which the proximal segment interacts with the front side of the C-domain (movie S1). This observation suggests that this orientation of the middle segment may exert a stabilizing effect; indeed, the corresponding clusters all map to low-temperature, low-energy segments in the unsorted trajectories (**Fig. S9**). Second, likely because of this stable association, the proximal segment can subsequently adjust its mode of engagement; for example, in movie S6, it shifts from interacting with the back groove to engaging the interface between the N- and C-domains, an engagement similar to the other back-side interactions shown in movies S2-S5.

Taken together, these observations suggest that anchoring of the middle segment is the more dominant element in tail engagement with the main body.

## DISCUSSION

The intrinsic disorder and large configurational freedom of the 62-residue βarr1 tail in its active state place it at the limits of what conventional structural analysis can meaningfully resolve. Enhanced sampling through TREMD substantially broadened the explored conformational space, but this increased heterogeneity also challenged geometric clustering approaches: RMSD- or linear projection-based methods failed to organize the ensemble into interpretable states without manual feature selection. By combining TREMD with a transformer-based autoencoder (TAE), we introduce a framework that shifts the analytical focus from predefined structural metrics to learned residue-level coupling patterns. In doing so, the present work positions attention-based representation learning not merely as a dimensionality reduction tool, but as a mechanism for extracting physically interpretable modular organization from highly heterogeneous MD ensembles. A key feature of this workflow is that attention-derived modules provide a residue-level framework for interpreting latent-space clusters, linking global conformational substates to segment-specific tail behavior (**Fig. 2**).

Many unsupervised learning approaches for molecular simulation data have focused on discovering slow collective variables, learning nonlinear reaction coordinates, or constructing kinetic models such as MSMs (reviewed in (Glielmo et al. 2021)). Autoencoder-based and variational formulations have been particularly useful for learning compact representations of molecular dynamics (Wehmeyer and Noe 2018). Related spectral methods for molecular kinetics, including TICA, VAMP, and EDMD, can extract slow dynamical modes, but their performance depends sensitively on the choice of observables used to represent the underlying dynamics (Ngo et al. 2023). However, these approaches generally emphasize global low-dimensional coordinates or discrete state assignments rather than residue-indexed relationships. In contrast, by treating each residue as a token, the self-attention mechanism in our TAE provides a residue-level view of how information from different tail residues is integrated by the model. These attention patterns allowed us to partition the flexible tail into interpretable residue groups without manually defining the segments in advance.

Although only tail residues are treated as tokens in the attention mechanism, each token is represented by its distance profile to the entire protein, so coordinated responses to main-body structure and dynamics are implicitly embedded in the learned features. Thus, the attention patterns reflect not only within-tail coupling but also how tail residues collectively respond to the structural context of the main body. This formulation enables simultaneous representation of internal conformational variability and overall rigid-body reorientation in intrinsically disordered or highly flexible protein regions by embedding both degrees of freedom within residue–distance patterns learned directly from the ensemble. Together, this approach reframes MD ensemble analysis as the task of learning cooperative residue organization from conformational ensemble, rather than defining states through expert-selected contacts, alignment schemes, or geometric thresholds.

A central finding is that the βarr1 tail exhibits a state-dependent modular architecture. In the basal state, the attention-derived segmentation recapitulates the known anchoring of the middle segment within the N-domain groove while resolving the proximal and distal regions as modules that are decoupled from each other. Consistent with this organization, the distal segment can sample alternative configurations within the extended N-domain groove without disrupting the anchored middle segment. In addition, the identification of a partially disengaged intermediate, in which the C-terminal portion of the middle segment releases from polar core interactions while other groove contacts persist, indicates that engagement with the N-domain groove and stabilization of the polar core are mechanistically separable events.

Upon activation and displacement of the tail from the N-domain groove by the receptor phosphopeptide, this modular organization reconfigures. In the active ensemble, attention outside the hinge region becomes comparatively weak and diffuse, whereas the hinge residues dominate the attention map. This redistribution reflects reduced residue–residue cooperativity as the tail becomes less constrained and more spatially dispersed. The hinge emerges as the principal orientational anchor, while the remainder of the tail samples a wide range of conformations with diminished persistent coupling. Despite this overall attenuation of cooperativity, the model resolves four segments beyond the hinge, including a subdivision of the distal region, indicating increased heterogeneity and altered coupling relationships. Notably, the distal-I/distal-II boundary is also broadly consistent with βarr isoform divergence in this region (**Fig. 7**): βarr2 lacks the βarr1 distal-II extension, whereas the C-terminal end of distal-I diverges between the two arrestins, suggesting that this attention-derived subdivision may represent an isoform-sensitive feature of the C-terminal tail organization. Latent-space clustering showed that segment-level variability in the active state arises from intertwined changes in internal tail geometry and orientation relative to the main body, two contributions that cannot be disentangled by a single alignment scheme.

The active-state ensembles reveal a pronounced bias toward positioning of the tail along the back side of the main body, together with multiple modes of occupation of the central crest crevice by the middle and/or distal segments. Although reminiscent of recent cryo-EM observations of a helical “chameleon motif” occupying this pocket in βarr2 (Maharana et al. 2024), our simulations suggest that crevice occupation encompasses several related configurations that vary in insertion depth and segmental contribution. Although rigorous kinetics cannot be extracted from the present TREMD data, inspection of representative trajectories suggests a qualitative ordering of engagement events. The proximal segment first associates with the main body, after which the middle segment approaches the central crest region. Once the middle segment anchors near the crevice, it remains associated while the proximal segment can still rearrange. This pattern is consistent with middle-segment anchoring as the more dominant determinant of sustained tail engagement with the main body.

Because the central crest region serves as the binding site for intracellular loop 2 (ICL2) of GPCRs during receptor core engagement, self-occupation of this site by the βarr1 tail would be expected to sterically hinder ICL2 insertion. This suggests an additional layer of autoinhibition: distinct from basal N-domain groove anchoring, crevice occupation would not block phosphopeptide binding but would selectively disfavor full core engagement, thereby favoring tail-only complexes and reducing the likelihood of transition to a core-engaged state. In this mechanistic framework, the tail is not merely released upon activation but acts as a regulator that biases arrestin toward alternative engagement modes through favorable self-occupation of a functionally critical surface.

More broadly, these findings provide a structural basis for how arrestin activation can encode signaling specificity. The tail’s modular and flexible nature indicates that it is not a binary switch, but a tunable reservoir of conformations. Distinct receptor phosphorylation patterns could differentially shape this conformational ensemble, thus exposing or hiding specific binding motifs for downstream effectors like clathrin or various kinases scaffolded by βarr (Lefkowitz and Shenoy 2005). Although the present analysis focuses on V2Rpp-bound βarr1, the framework is readily extendable to other receptor phosphopeptides and to βarr2, enabling systematic comparison of isoform- and receptor-specific tail dynamics, and more broadly to proteins containing long, flexible regions sampled by TREMD.

## METHODS

The temperature replica-exchange molecular dynamics (TREMD) simulation and analysis protocols for the basal and V2Rpp-bound active state of βarr1 were described in the companion manuscript (Ngo et al. 2025).

### Data preparation for transformer-based autoencoder (TAE) modeling

In this study, the TAE models treated each residue as a token, represented by a feature vector of its distances to all other residues. To construct the training dataset, we used the temperature-sorted TREMD replicas 35–46, spanning 306.9 to 312.4 K, to enrich sampling of conformations near the physiological temperature of 310 K. The frames from these replicas were processed through a custom pipeline that automated loading, normalization, and organization of residue-level representations, ensuring consistent formatting across replicas for subsequent TAE model training and analysis.

Specifically, the coordinates of all Cα atoms were retrieved with the VMD Python API, and for each frame, the pairwise distance matrix among all Cα atoms was computed as *D_ij_* =‖ *r_i_* – *r_j_* ‖_2_, yielding a three-dimensional tensor of shape (frames, residues, residues). Each feature, corresponding to the distance from a token residue to another residue, was standardized by z-scoring across all frames and token residues using StandardScaler. For analyses focused on the βarr1 tail, the token set was restricted to the tail residues, while each token retained its distance profile to all Cα atoms in the protein.

To ensure data quality, frames exhibiting large consecutive Cα-Cα distance (>4.5 Å), indicative of the βarr1 tail crossing a periodic boundary even after re-centering of the center of mass (COM) of the main body, were identified and excluded to eliminate potential boundary artifacts. However, for the low-temperature TREMD replicas (replicas 25-49; 302.0-313.9 K), no such frames were detected.

Our framework can aggregate multiple trajectories across user-specified replica indices or frame ranges, producing a unified dataset with consistent feature dimensions and indexing. Each residue token thereby corresponds to a distance profile relative to all other residues, allowing residue-wise modeling of the conformational space explored by the tail.

### TAE architecture

Building on the residue token representation defined above, a TAE framework was developed with PyTorch (v2.8.0) to learn low-dimensional representations of residue-residue distance patterns from TREMD ensembles. For each frame, the input to the model was a matrix whose rows corresponded to residue tokens and whose columns represented their distance features. The transformer encoder first projected each token through a linear layer into a 128-dimensional embedding space (defined as the model dimension, d_model) and added learnable positional embeddings to provide positional context for each residue. It then applied a stack of four self-attention layers, each combining multi-head (four heads) self-attention, a two-layer feed-forward transformation, and post-layer normalization, allowing the model to capture both local and long-range dependencies among residues. During the forward pass, the model retained the attention weights to enable downstream analysis of attention patterns.

Three alternative readout strategies were implemented and compared: i) mean pooling, which averages over token embeddings; ii) a classification (CLS) token readout, in which a special learnable token is prepended to the input and trained to summarize global information about the entire tail; and iii) Pooling by Multi-head Attention (PMA), which employs a small number of learnable seed queries (r) to produce permutation-invariant summaries of the residue set. We used the PMA readout with r = 3 for the basal-state models and r = 5 for the active-state models, which provided the strongest attention signaling in the corresponding βarr1 tail analysis (see Results). These PMA slot vectors were then combined into a single summary by a slot-mixing multi-head attention block, in which queries were derived from a learnable window token and keys and values were derived from the PMA slots.

A latent bottleneck layer compressed the encoder readout into a 64-dimensional embedding, which was subsequently expanded back to d_model and used as the decoder memory. The transformer decoder consisted of multi-head attention and feed-forward blocks, architecturally mirroring the encoder. The model reconstructed the residue–residue distance profiles from the decoder outputs using a final linear projection and was trained by minimizing the mean squared reconstruction loss. This bottlenecked reconstruction objective encouraged the latent representation to capture the most informative residue–residue distance relationships.

### Training and validation of TAE models

Model training and validation procedures were designed to ensure stable optimization, appropriate weighting of structural features, and reproducible evaluation across ensembles.

During training, a sequence-separation-dependent weighting scheme was applied so that the reconstruction of short-range contacts and distances involving the hinge residues contributed less to the reconstruction loss. This weighting emphasized medium- and long-range couplings that characterize collective tail rearrangements, while down-weighting both local fluctuations and the dominant influence of the hinge. Specifically, a sequence-separation dependent weight matrix W was constructed to modulate the reconstruction objective. This rectangular matrix of shape T×R (tokens × residues) assigned each element W_ij_ according to the sequence separation |I_i_ − J_j_| between the token residue I_i_ and feature residue J_j_. Specifically, a ramp function {w(0)=0.0, w(1)=0.0, w(2)=0.1, w(3)=0.3, w(4)=0.5, w(d≥5)=1.0} was applied to this matrix so that near-neighbor pairs contributed less to the mean-squared reconstruction loss. To further reduce the influence of the first five residues connecting the tail to the main body, the corresponding token rows of W were downweighted by factors (Levin et al. 2021 0.7, 0.9), respectively. Finally, W was normalized to have approximately unit mean before use in the loss function. This weighting scheme ensures that training focused on medium- and long-range residue conformations underlying the collective motions of the flexible tail. The first few residues were nonetheless retained as tokens to capture how hinge dynamics modulate tail flexibility.

For the basal-state TAE models, training was performed using the Adam optimizer with an initial learning rate of 0.0004 and a batch size of 256 for up to 200 epochs, with a multi-stage cosine-annealing schedule and early stopping based on the validation loss (patience 15). For the active-state TAE models, training followed the same scheduling and early-stopping criteria but used an initial learning rate of 0.0005 and a batch size of 512, reflecting both the greater conformational heterogeneity of the active ensemble and the larger available dataset size. While larger batch sizes generally improve clustering performance, batch size selection was constrained by dataset size to ensure an adequate number of batches per epoch for stable training. The observation that larger batch sizes yield better clustering results (e.g., higher silhouette and DBCV scores) despite slightly higher validation loss is consistent with established principles in representation learning. Although validation loss primarily reflects per-sample reconstruction fidelity, it does not necessarily correlate with the quality or geometric structure of the learned latent space. Studies on optimization dynamics show that small-batch training introduces higher gradient noise, which, while often beneficial for finding flat minima and improving generalization, can result in scattered or distorted latent space geometry. Conversely, larger batch sizes yield lower-noise, more stable gradient estimates, leading to smoother optimization trajectories and more cohesive latent space organization that better supports downstream clustering tasks (Keskar et al. 2016). Therefore, the improved clustering scores with larger batches reflect enhanced latent-space geometric quality rather than superior reconstruction fidelity.

Although the twelve temperature-sorted replicas formally represent slightly different thermodynamic ensembles, their temperatures lie within a narrow window near 310 K, and we therefore treated them as an approximate single ensemble for training. With this approximation, shuffling was enabled during training, allowing each batch to draw frames from across all replicas and avoiding artifacts associated with locally correlated frames while yielding reconstruction and clustering results superior to unshuffled training. The reconstruction loss served as the training criterion, and gradients were clipped to a maximum L2 norm of 0.2 after an initial 25-epoch burn-in period to prevent rare gradient spikes.

During model training, deterministic random seeds were used to ensure reproducible data loading and optimization behavior. Replica trajectories were drawn from a fixed set of 12 temperature-sorted replicas (35–46), which were partitioned deterministically into training and validation sets by enumerating all possible 10–2 splits. This procedure yielded 66 distinct train–validation combinations, and a separate TAE model was trained for each split, providing a full ensemble of models for downstream attention averaging and module identification. For clustering analyses, the model achieving the lowest reconstruction loss on the tail region was selected as the representative model.

### Attention analysis and module identification

After training, attention maps were generated for each TAE by running the TREMD dataset through the model and storing the attention weights from all heads and layers. For each model, attention weights were first normalized so that, for every query residue (token), the attention values it assigned to all tokens (excluding the CLS token if present) summed to one. The normalized weights were then averaged over all batches, heads, and layers to obtain a single residue–residue attention matrix for that model. To obtain an ensemble-averaged representation, these model-level attention matrices were further averaged across all independently trained TAE models for a given condition (basal or active).

To identify structural modules within the βarr1 tail, hierarchical agglomerative clustering with average linkage was applied to precomputed distance matrices derived from the attention patterns. The distance between residues was defined using a profile-based cosine distance computed on the concatenated row-normalized query-key profiles [A | A^T^], where A denotes the ensemble-averaged attention matrix.

### Frame embedding and clustering analysis

To select the best model for downstream analysis, reconstruction quality was evaluated for each trained TAE model using simulation frames from sorted temperature replicas 37–46 (near 310 K). Residue-distance features were computed and normalized as described above, then passed through the full autoencoder. Mean-squared error loss was computed over tail-to-tail and tail-to-body regions on a subset of 20 batches, and the model achieving the lowest reconstruction loss on the tail-to-tail block was selected. Using this model’s encoder, latent embeddings were then extracted for all frames to obtain low-dimensional representations of tail conformations. These embeddings were stored with corresponding frame and trajectory metadata for downstream clustering.

To identify conformational states across the ensemble, the 64-dimensional latent embeddings were first L2-normalized to unit length and then subjected to a two-stage dimensionality reduction and clustering workflow combining Uniform Manifold Approximation and Projection (UMAP) and Hierarchical Density-Based Spatial Clustering of Applications with Noise (HDBSCAN) algorithms. A nested grid search systematically explored UMAP parameters (n_components, n_neighbors, min_distance, and metric) together with HDBSCAN parameters (min_cluster_size, min_samples, and metric). For each configuration, clustering quality was assessed using silhouette, modified silhouette (excluding noise points), and density-based cluster validity (DBCV) scores. Configurations that produced at least five clusters, a silhouette score above 0.30, and a noise fraction below 15% were retained. Among these, the parameter set yielding the highest DBCV score was selected as optimal (Tables S2 and S3).

Using the optimal parameters from the grid search, clustering was further refined by testing principal component analysis (PCA) preprocessing at three variance retention thresholds (0.90, 0.95, and 0.99) to reduce noise. When PCA preprocessing improved the DBCV score, the PCA-transformed embeddings were used for final clustering.

Using the best-performing UMAP embedding, K-means clustering was performed with cluster numbers spanning n ± 15, where n represents the HDBSCAN cluster count. HDBSCAN preferentially identifies variable-density structure in irregular or noisy data, whereas K-means optimizes within-cluster variance and is best suited to compact, approximately spherical clusters under Euclidean geometry.

Direct application of K-means to the original latent embeddings, with or without PCA preprocessing, yielded consistently low silhouette scores (<0.12 for both basal- and active-state models), indicating weak centroid-separable structure in the latent space. In contrast, K-means applied to the UMAP-derived embeddings produced substantially higher silhouette values (>0.75 for both models), demonstrating markedly improved geometric separation following nonlinear dimensionality reduction that preserves local neighborhood structure. Thus, K-means clustering was conducted in the UMAP embedding space, where cluster structure was clearly resolved and supported by internal validation metrics.

To assess whether the observed cluster organization depended on the choice of clustering algorithm, the final HDBSCAN partitions were compared with K-means solutions obtained in the same UMAP space (**Fig. S4**). The two methods produced largely concordant partitions, supporting the presence of well-resolved conformational basins in the UMAP embedding rather than assignments imposed by a particular clustering algorithm. Cases of partial disagreement were used as diagnostic information, as they typically reflected elongated or orientation-dispersed regions of the embedding that were density-connected under HDBSCAN but not well represented by the compact, centroid-based partitions produced by K-means.

## Author Contribution

MR, VN, JAJ, LS conceptualized the work. MR and LS designed and carried out the computations and developed the analysis approaches. MR, VN, JAJ, and LS analyzed the data. MR and LS wrote the initial draft, and all authors contributed to finalizing the manuscript.

## Acknowledgements

Support for this research was provided by the National Institute on Drug Abuse–Intramural Research Program, Z1A DA000606 (L.S.), by NIMH R01 MH541397 (J.A.J.) and the Hope for Depression Research Foundation (J.A.J.), and by the Advanced Scientific Computing Research (V.N.), the Office of Science at DOE (No. ERKJZN1). This research used resources of the Oak Ridge Leadership Computing Facility (OLCF) at the Oak Ridge National Laboratory, which is supported by the Office of Science of the U.S. Department of Energy under Contract No. DE-AC05-00OR22725. The TREMD simulations were carried out on the Frontier supercomputer at the OLCF, under the INCITE 2024 Award BIP248. The TAE trainings were performed using the NIH high-performance computing Biowulf cluster (https://hpc.nih.gov). Data analysis utilized computational resources on both the Andes cluster at OLCF and Biowulf.

