## Supplemental Information for "Transformer-based framework uncovers state-dependent modular organization and conformational landscapes of the β-arrestin 1 C-terminal tail"

### SUPPLEMENTAL RESULTS

#### Assessment of clustering resolution and selection of the final active-state parameters

To assess whether the relatively large number of clusters identified in the active ensemble reflects genuine structural heterogeneity rather than over-fragmentation, we evaluated clustering behavior along a local HDBSCAN parameter path while holding the UMAP embedding fixed and enforcing a minimum cluster size of 30 frames to avoid interpreting isolated, sparsely sampled microclusters (**Table S2**).

Within this constrained parameter space, increasing hdbscan_min_samples from 15 to 25 led to a monotonic improvement in internal clustering metrics. The density-based cluster validity (DBCV) score increased from 0.597 to 0.614 to 0.627, and the modified silhouette score increased from 0.798 to 0.808 to 0.819, as min_samples increased from 15 to 20 to 25. Over the same range, the number of clusters decreased only modestly, from 782 to 765 to 738, while the noise fraction remained moderate, increasing from 0.112 to 0.119 to 0.123. Thus, tightening the density criterion improved cluster quality without causing a qualitative change in clustering resolution. The persistence of several hundred clusters across this parameter range supports that the high cluster count is a robust feature of the embedding rather than a consequence of an overly permissive clustering regime.

Further increasing min_samples to 30 made the density criterion too restrictive, causing the noise fraction to exceed the predefined acceptance threshold (>0.135). We therefore selected hdbscan_min_samples = 25 as the most stringent density setting that still preserved acceptable data coverage while giving the strongest cluster-validity metrics within the prespecified minimum-cluster-size constraint.

To further assess whether the resulting large number of clusters reflects over-fragmentation, we examined the distribution of cluster sizes for the selected configuration. The cluster-size spectrum shows a smooth, approximately heavy-tailed decay without a pronounced elbow or an accumulation of clusters at the minimum allowed size (**Fig. S4**). Because the smallest retained clusters contain approximately 30 frames, consistent with the imposed lower bound, the absence of a strong pile-up at this boundary argues against a solution dominated by artificially truncated microclusters. Instead, cluster sizes decrease gradually across ranks, consistent with a heterogeneous conformational landscape with many metastable states spanning a range of populations.

Together, these analyses indicate that the final clustering solution is stable and well supported by the data. Within the parameter regime defined by a minimum cluster size of 30 frames, increasing the density threshold improved cluster validity while only gradually reducing the number of recovered clusters, and the selected solution maintained acceptable noise coverage without evidence of excessive fragmentation. We therefore conclude that the approximately 700-cluster decomposition used for the final analysis reflects intrinsic conformational heterogeneity of the active ensemble rather than parameter-driven over-clustering.

### SUPPLEMENTAL TABLES

#### Table S1. Summary of TREMD simulations

Each REMD simulation was carried out using 200 replicas, employing the 290 – 400 K temperature ladder described in (Ngo et al. 2025).

| state | Length/replica (ns) |
|---|---|
| basal | 1122 |
| v2rpp-bound | 3800 |

**Table S2. Representative UMAP-HDBSCAN grid-search parameters and clustering performance metrics for the active-state ensemble**

The entries shown represent a selected subset of a large joint UMAP–HDBSCAN parameter screen. For this subset, HDBSCAN was applied to the UMAP embedding of the active-state TAE latent space generated with n_components = 3, n_neighbors = 15, min_dist = 0.0, and metric = correlation. The subset spans a local HDBSCAN parameter grid varying minimum cluster size (min_cluster_size = 25 or 30), density threshold (min_samples = 15, 20, 25, or 30), and distance metric (euclidean or manhattan), yielding the 16 representative parameter combinations shown. The parameter set selected for downstream analyses, min_cluster_size = 30, min_samples = 25, and metric = euclidean, is highlighted in yellow. This configuration represented the most stringent density setting that maintained acceptable data coverage, defined as a noise fraction ≤ 0.135; for parameter sets exceeding this noise-fraction threshold, DBCV was not calculated. Among the displayed parameter combinations satisfying this coverage criterion, the selected configuration produced the strongest cluster-validity scores, including DBCV and modified silhouette, within the prespecified minimum-cluster-size constraint.

| HDBSCAN | | | | | | silhouette | modified silhouette | DBCV |
|---|---|---|---|---|---|---|---|---|
| min cluster size | min samples | metric | clusters | number of noise | noise ratio | | | |
| 30 | 15 | euclidean | 782 | 7399 | 0.112 | 0.605 | 0.798 | 0.597 |
| 30 | 15 | manhattan | 786 | 7789 | 0.118 | 0.596 | 0.799 | 0.597 |
| 30 | 20 | euclidean | 765 | 7859 | 0.119 | 0.602 | 0.808 | 0.614 |
| 30 | 20 | manhattan | 754 | 7836 | 0.119 | 0.603 | 0.808 | 0.608 |
| 30 | 25 | euclidean | 738 | 8090 | 0.123 | 0.606 | 0.819 | 0.627 |
| 30 | 25 | manhattan | 740 | 8491 | 0.129 | 0.597 | 0.821 | 0.620 |
| 30 | 30 | euclidean | 710 | 9001 | 0.137 | 0.596 | 0.835 | |
| 30 | 30 | manhattan | 704 | 9216 | 0.140 | 0.593 | 0.837 | |

### SUPPLEMENTAL FIGURES AND LEGENDS

**Figure S2. Model-to-model variability of attention matrices in basal- and active-state TAE ensembles.**

Panels A and B show the basal-state attention variability, and panels C and D show the active-state attention variability. Panels A and C show pixel-wise standard deviations (SDs) of residue–residue attention values across independently trained TAE models. Panels B and D show the corresponding coefficient-of-variation (CV) maps, calculated as SD divided by the ensemble-averaged attention value. In all panels, rows represent query residues and columns represent key residues, with both axes corresponding to βarr1 tail residues 357–418.

In both basal and active ensembles, the SD maps show that model-to-model variability is concentrated mainly in regions that also carry strong mean attention, rather than appearing as unrelated off-pattern noise. The CV maps further indicate that the dominant attention features exhibit moderate relative variability across models, supporting the robustness of the overall module-level organization. However, local variation in attention strength remains, particularly near high-attention regions and module boundaries.

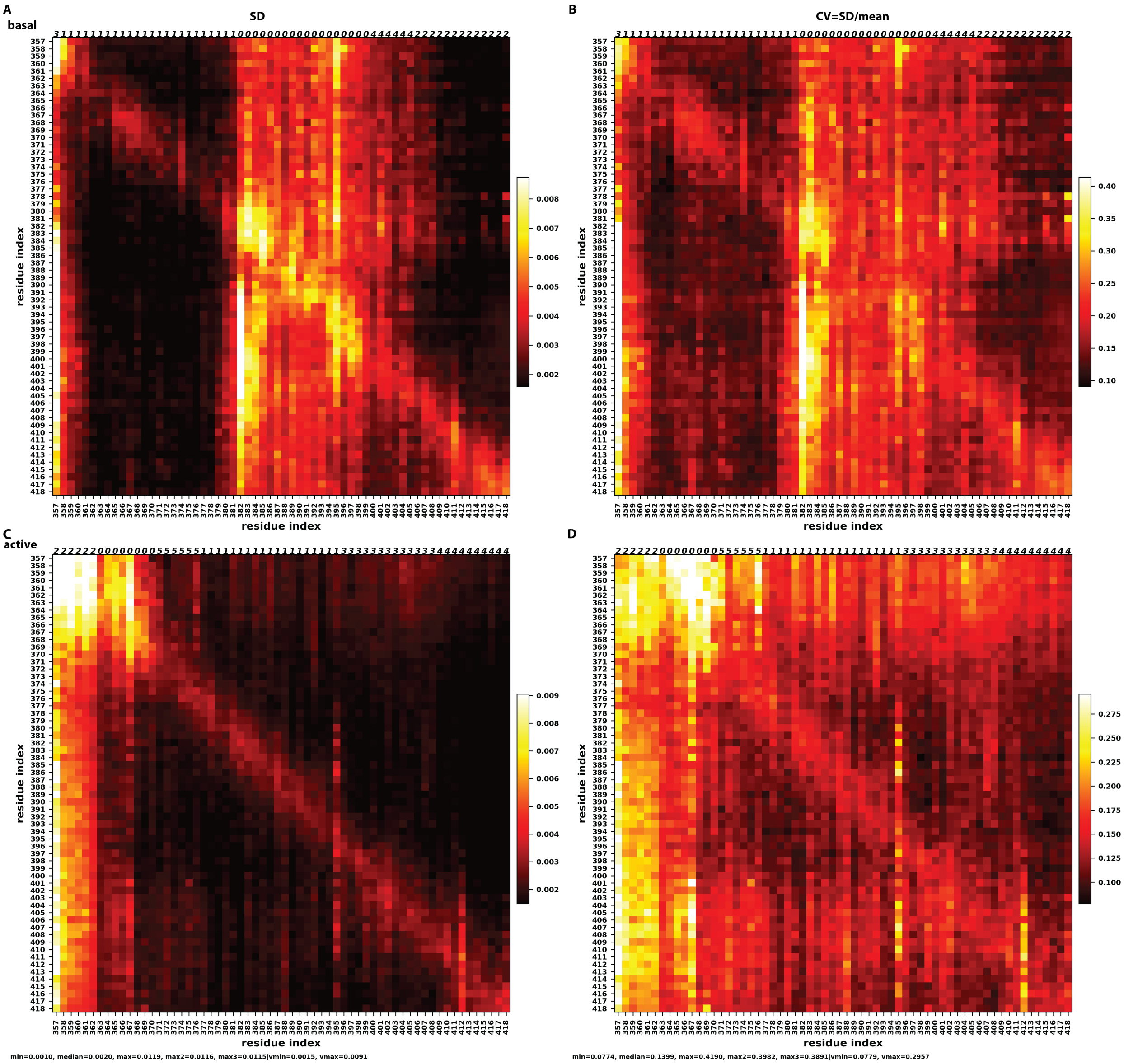
A
basal
SD
B
CV=SD/mean
C
active
D
residue index
min=0.0010, median=0.0020, max=0.0119, max2=0.0116, max3=0.0115|vmin=0.0015, vmax=0.0091
min=0.0774, median=0.1399, max=0.4190, max2=0.3982, max3=0.3891|vmin=0.0779, vmax=0.2957

**Figure S3. Averaged attention matrices of the 57-token models of βarr1 active ensembles displayed similarity to the 62-token models.**

The ensemble-averaged attention matrix for the 57-token active-state ensemble was generated using the same model-averaging and clustering pipeline used for the 62-token models and exhibited similar attention patterns and modular organization. To preserve a minimum segment length of at least 10 residues, clusters 3, 5, and 2 from the clustering solution were merged into the proximal segment.

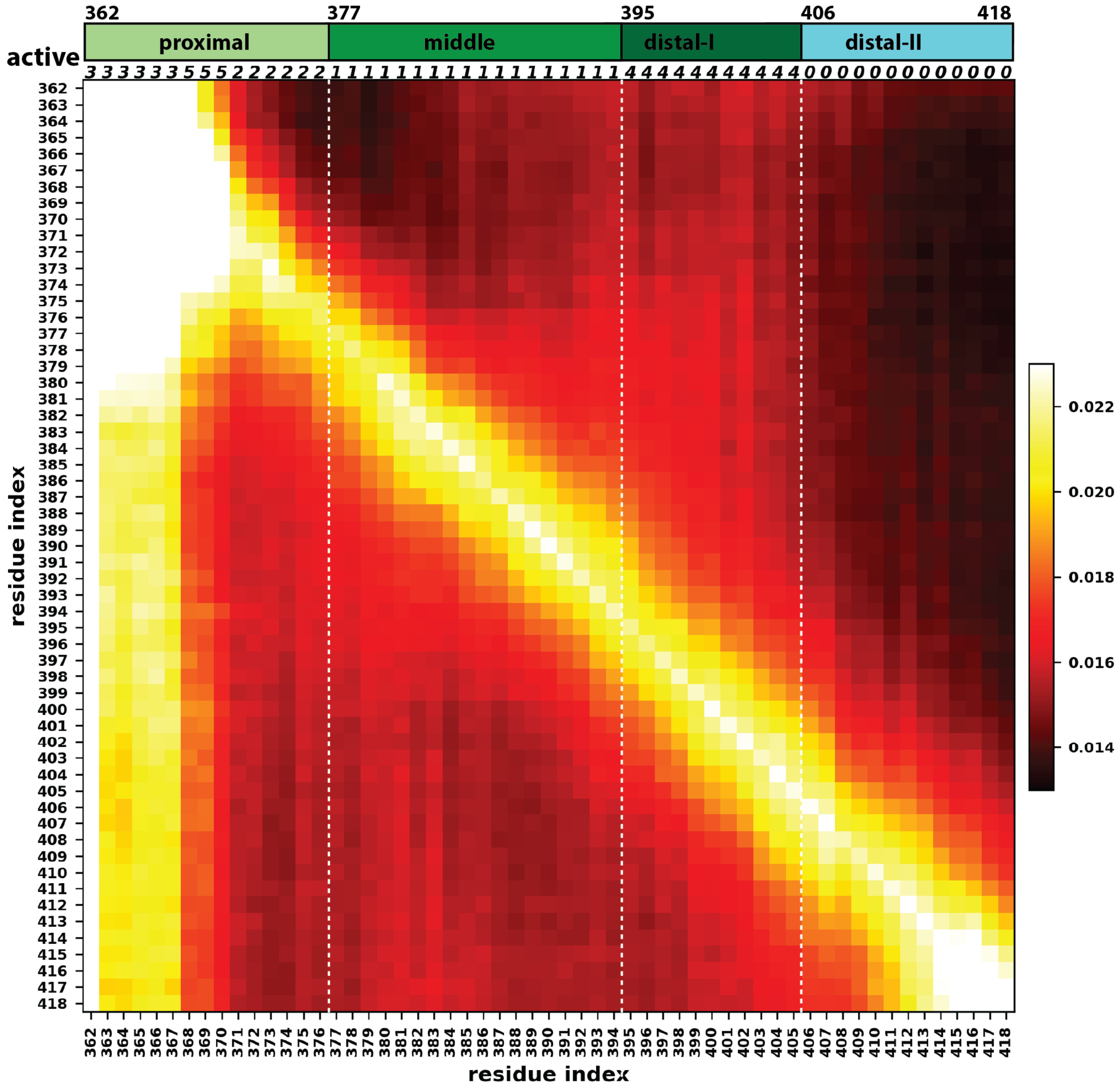

**Figure S4. Comparison of cluster-size spectra from HDBSCAN and K-means clustering of the active-state ensemble**

Comparison of cluster population distributions obtained from HDBSCAN (blue) and K-means (red) clustering of the TAE latent embedding for the active-state ensemble. Clusters are ordered from largest to smallest by frame count; for HDBSCAN, noise points (unassigned frames) are excluded from the ranking and statistics. The y-axis shows the number of frames assigned to each cluster. Discrete cluster sizes are shown as smoothed curves using cubic splines to illustrate the decay in cluster population across ranks.

The two methods yield clearly different cluster-size spectra. HDBSCAN shows a much larger top-ranked cluster followed by a broader heavy-tailed decay, whereas K-means distributes frames more evenly across much of the rank range and yields progressively smaller clusters at higher ranks. This difference is consistent with the underlying logic of the two methods: K-means enforces a complete centroid-based partition of the embedding, whereas HDBSCAN identifies density-connected regions and leaves low-density boundary points unassigned as noise. In the present context, the HDBSCAN distribution likely provides a more meaningful description of the landscape, as it captures an uneven population structure with several highly populated states and many progressively less populated states.

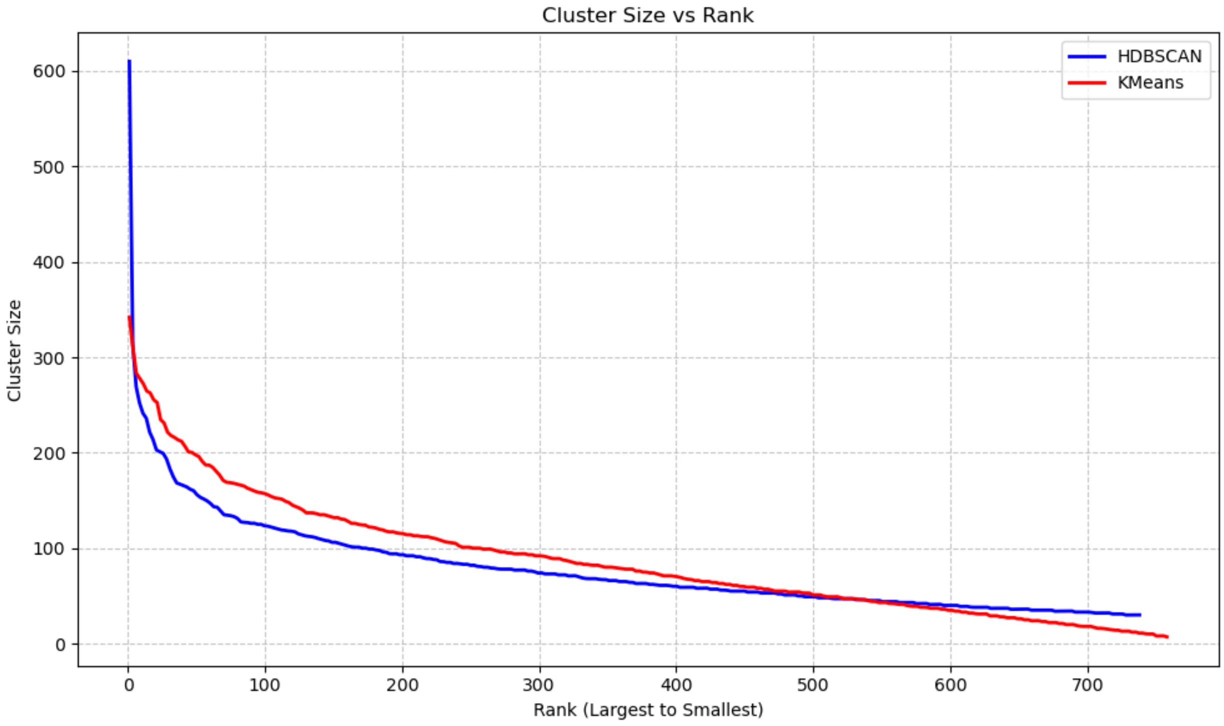
Cluster Size vs Rank
HDBSCAN
KMeans
Cluster Size
Rank (Largest to Smallest)
600
500
400
300
200
100
0
0
100
200
300
400
500
600
700

**Figure S5. Selected active-state cluster conformations illustrating segmental decoupling of tail organization.**

Panels A shows that in cluster a7, the proximal and middle regions are comparatively well defined, while the distal-II region is substantially more dispersed. In cluster a9 (B), the proximal region is relatively well defined, whereas the middle, distal-I, and distal-II segments are markedly more variable. Panels C and D show clusters a14 and a35, respectively, which exhibit low main-body–tail RMSDs in the distal-II segment despite comparatively dynamic middle segments. Together, these examples illustrate that, as in the basal state, active-state clusters can display pronounced decoupling among tail segments, with relatively ordered conformations in some regions coexisting with substantial mobility in others. The colors of the main-body domains, V2Rpp, and each tail segment are indicated by the bars at the bottom.

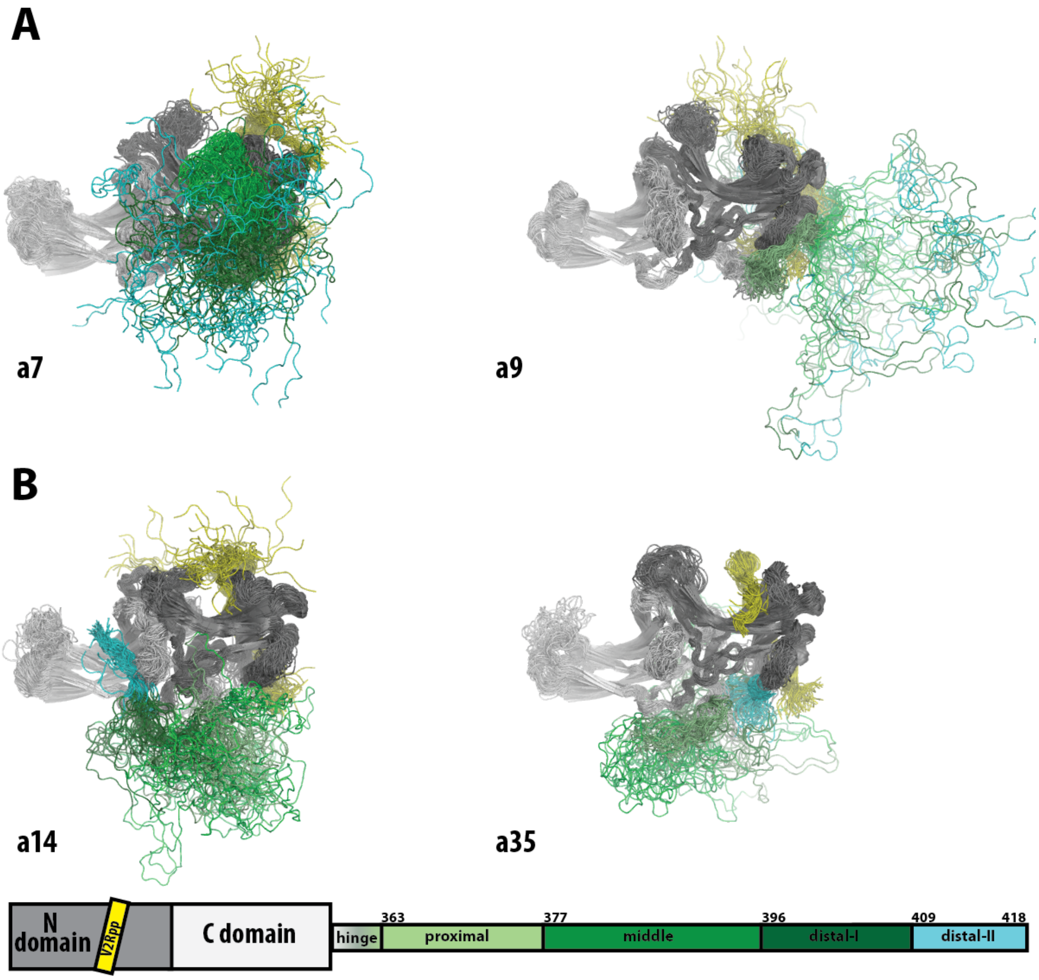

**Figure S6. Active-state clusters with large main-body–tail RMSDs but low tail–tail RMSDs.**

Panels A, C, E, and G show zoomed-out views of clusters a20, a40, a11, and a95, respectively, with frames superimposed on the βarr1 main body to illustrate the relative positioning, orientations, and spatial distributions of the tail. Panels B, D, F, and H show zoomed-in views of the tail segments that are relatively well defined within each cluster, with frames superimposed on the segment(s) with low tail-tail RMSDs (see **Fig. 4B**). Specifically, in panel B, cluster a20 is superimposed on the proximal and middle segments, revealing a defined local structure in which residues 388-395 form two helical turns. In panel D, cluster a40 is superimposed on the middle segment, showing a conformation similar to that of cluster a20 for the middle segment. In panels F and H, clusters a11 and a95 are superimposed on the distal-I and distal-II segments. The colors of the main-body domains, V2Rpp, and each tail segment are indicated by the bars at the bottom.

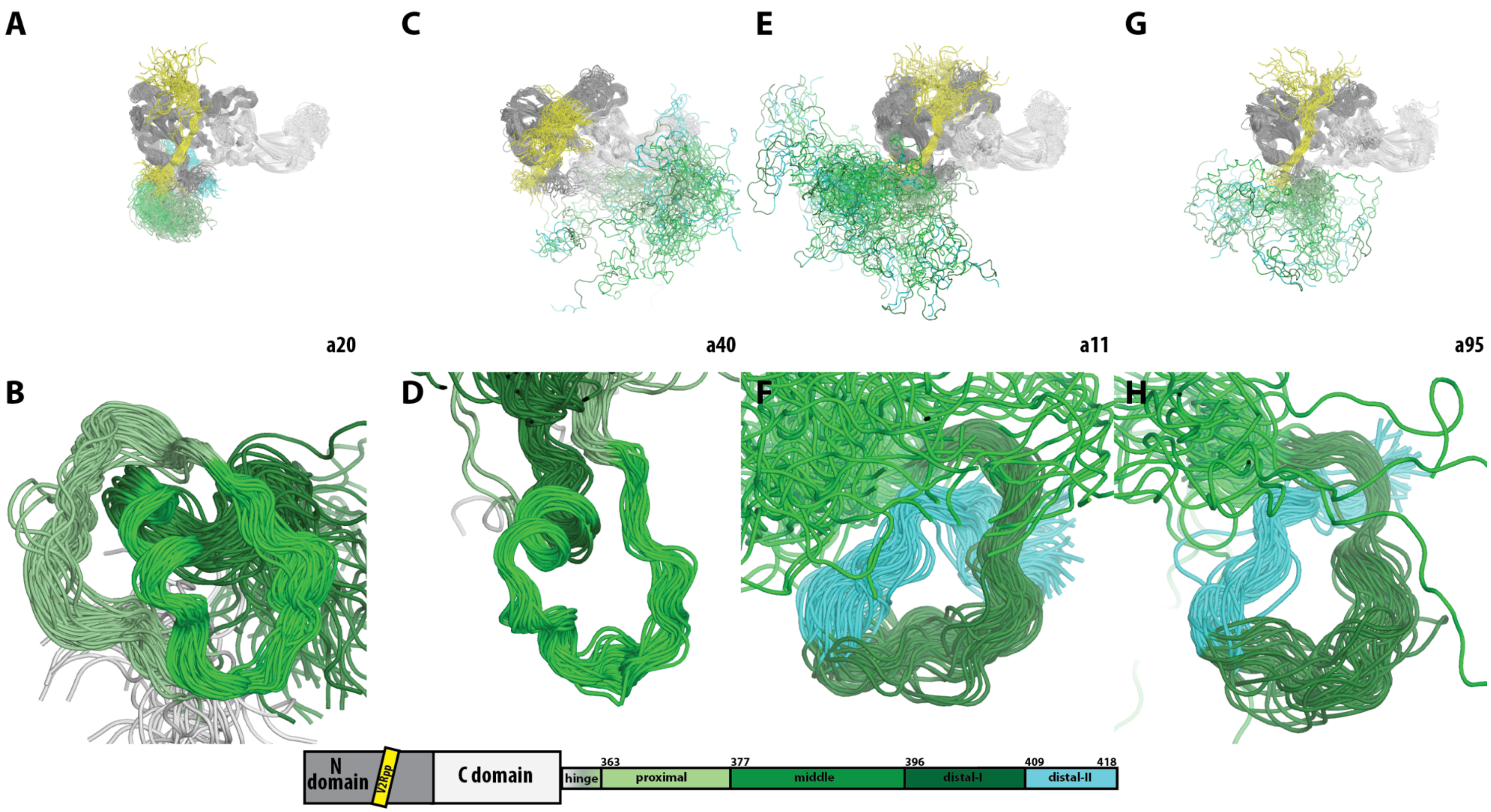
A
C
E
G
a20
a40
a11
a95
B
D
F
H
N domain
V2Rpp
C domain
363
377
396
409
418
hinge
proximal
middle
distal-I
distal-II

**Figure S7. Distance-based identification of back-side versus front-side tail engagement.**

Mean distances for residue pairs 374-307 and 375-255 are shown as scatter plots across HDBSCAN clusters, with 374-307 distance on the x-axis and 375-255 distance on the y-axis. Each point represents one cluster, and error bars indicate the pooled standard error across replicas. Panel A includes all clusters in which at least one of the two plotted distances is below 22 Å, as marked by the gray dotted lines, and provides an overview of the spatial relationship between these two indicators of tail positioning. Panel B shows the subset of clusters that additionally satisfy a third distance criterion, 392-244 < 17 Å, thereby isolating clusters with proximity at this third site and allowing comparison of how this additional constraint relates to the 374-307 and 375-255 distance distributions. Visual inspection of these 12 clusters confirmed that this distance threshold accurately captured clusters in which the βarr1 tail approached or entered the crevice (**Fig. 6**). The number of clusters in each panel is indicated by n, and cluster IDs are labeled next to the corresponding data points.

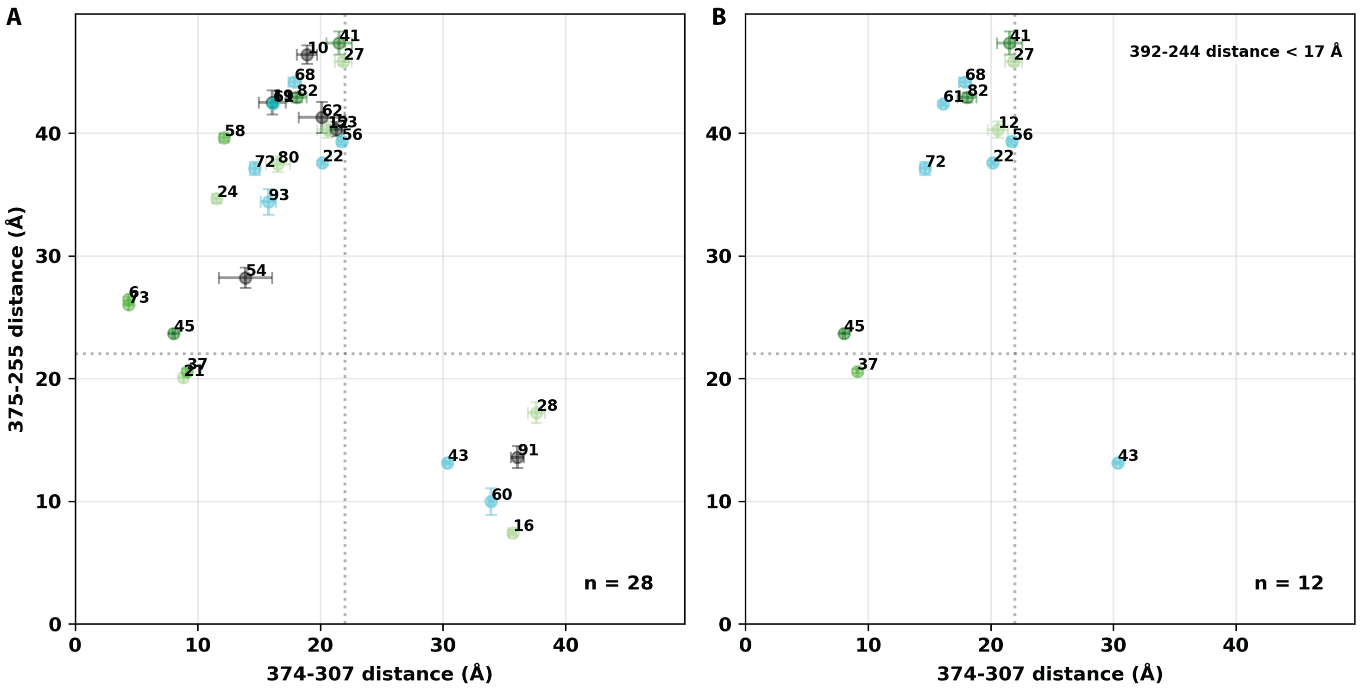

A
B
375-255 distance (Å)
374-307 distance (Å)
392-244 distance < 17 Å
n = 28
n = 12
41
10
27
68
82
62
56
58
72
80
22
24
93
54
6
73
45
37
21
28
43
91
60
16
61
12

**Figure S8. Clusters a22 and a56 share similar tail conformations but differ in main-body conformation.**

Superposition of clusters a22 and a56 shows that, despite their nearly identical tail conformations, they are associated with distinct main-body conformations. Panel A showed quantitative PIA analysis (Ngo et al. 2025) comparing the two clusters, which identified the most apparent differences around ASwI and α-helix I within the three-element interaction region associated with V2Rpp binding. Conformational rearrangements of these regions from cluster a56 to a22 are indicated by arrows in Panel B. These regions are shown in red for cluster a22 and blue for cluster a56. The N and C domains are shown in dark and light grey, respectively. These differences within the three-element interaction region likely result from allosteric modulation by distinct rearrangements of the finger loop (FL) and C-loop (CL) of the central crest crevice, in response to tail engagement near the crevice in the two clusters, as shown in panel A. Thus, these observations suggest an allosteric pathway between these functionally critical regions.

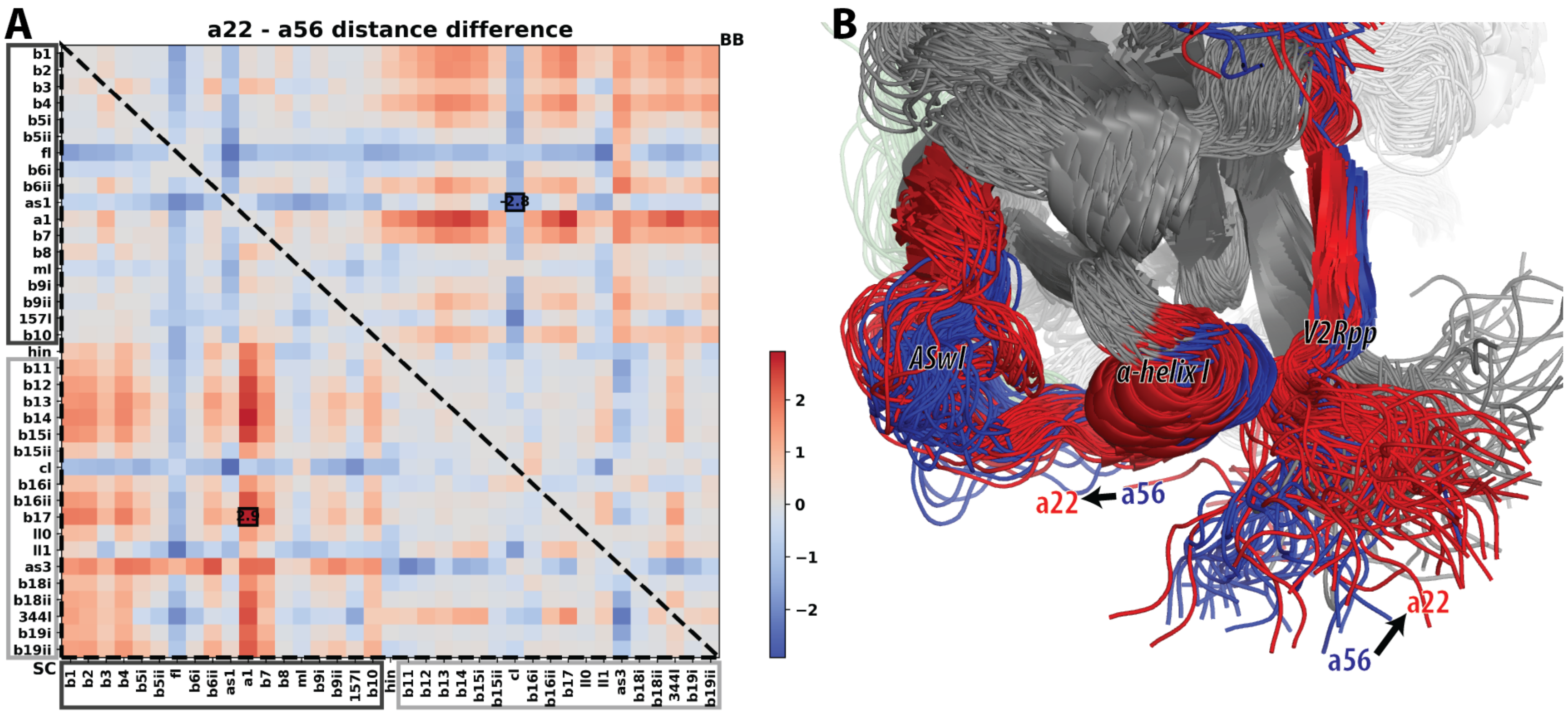
A
a22 - a56 distance difference
BB
SC
B
ASwI
a-helix I
V2Rpp
a22
a56
a22
a56

**Figure S9. Temperature and energy histories of six unsorted TREMD trajectories containing back-to-front middle-segment engagement with the central crest crevice.**

For each unsorted trajectory, with the trajectory index indicated at the top right, the temperature trace is shown on the left and the corresponding total energy trace on the right as a function of TREMD time. In the energy plots, the pink trace shows the raw energy time series and the black trace shows the smoothed profile. Blue horizontal bars mark the time intervals corresponding to the indicated TAE clusters. In all six cases, the frames corresponding to these clusters occur in relatively low-temperature, low-energy segments of the unsorted trajectories, consistent with stabilization after the middle segment engages the central crest crevice in a back-to-front orientation.

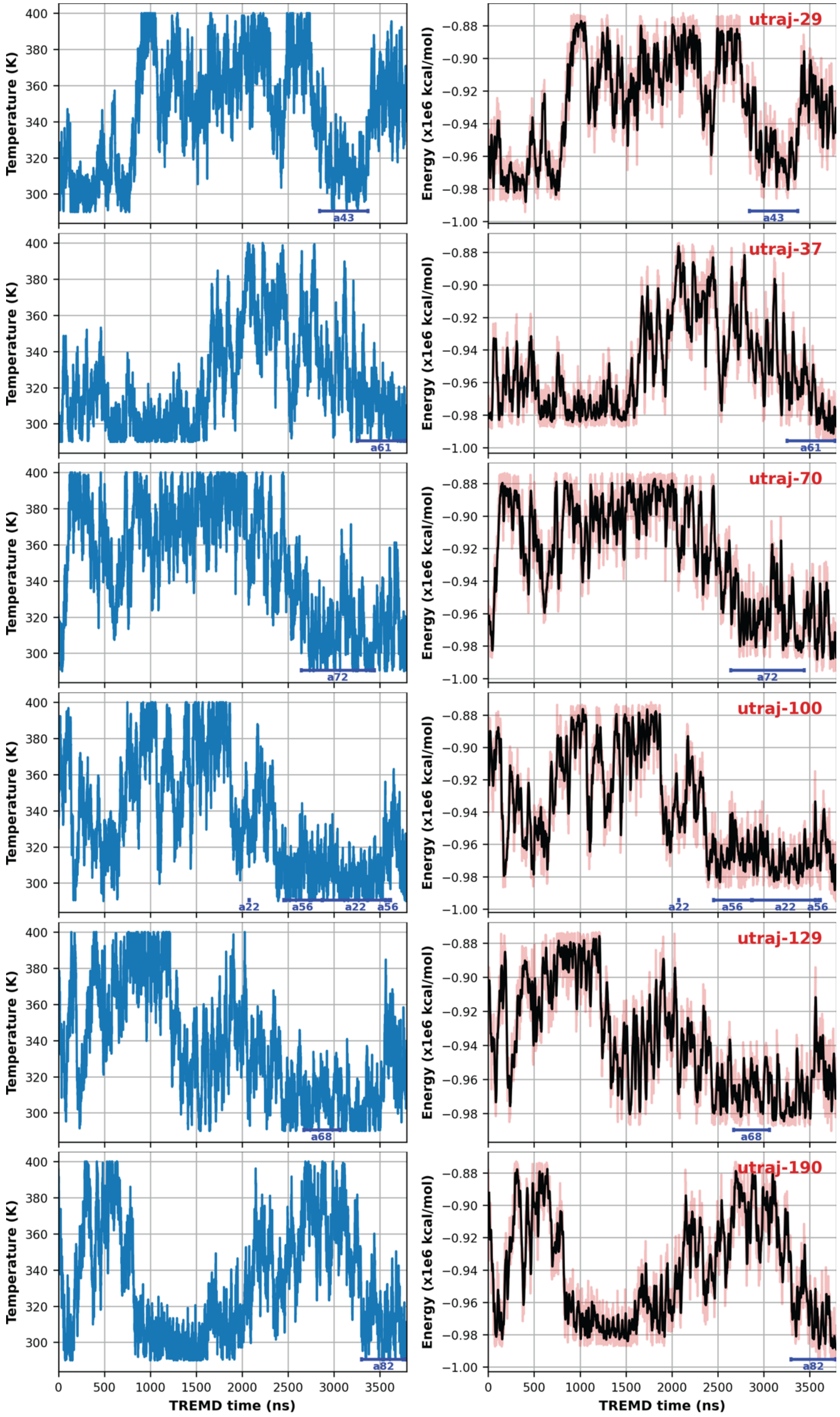
utraj-29
utraj-37
utraj-70
utraj-100
utraj-129
utraj-190
Temperature (K)
Energy (x1e6 kcal/mol)
TREMD time (ns)
a43
a61
a72
a22
a56
a22
a56
a68
a82
